# Synaptic and circuit mechanisms prevent detrimentally precise correlation in developing visual system

**DOI:** 10.1101/2022.11.11.516093

**Authors:** Ruben A. Tikidji-Hamburyan, Gubbi Govindaiah, William Guido, Matthew T. Colonnese

## Abstract

During development, retinal axons create broad and imprecise connections in the thalamus. This topology, very different from adults, supplies developing thalamic neurons with locally homogeneous synaptic currents and should cause spike correlation between thalamocortical neurons on a millisecond timescale. Such correlations have not been observed *in vivo*, at these ages, and would likely be maladaptive. Here, we use a biophysical model of the visual thalamus with the membrane and synaptic properties of 7-10 day-old mice to show that the developmentally appropriate dominance of NMDA-receptor currents and absence of strong recurrent inhibitory and excitatory connections prevents precise correlation and preserves topographic information in thalamic spikes. We illustrate possible reasons for this desynchronization using a phenomenological cortical model, which shows impaired network diversity when driven with precisely correlated inputs. Our results suggest that developing synapses and circuits evolved mechanisms to compensate for detrimental, “parasitic” correlation arising from the unrefined and immature circuit.

## Introduction

Correlated neural activity plays an essential role in circuit development and plasticity (***Katz and Shatz, 1996***). Gunther Stent, following Hebb’s hypothesis, proposed that synchronized activity among presynaptic neurons allows them to effectively depolarize and fire the postsynaptic cell to maintain their synapses, while neurons that are not coordinated lose their connections (***Stent, 1973***). Mechanisms that generate synchronous spontaneous activity during initial circuit formation have been identified in every brain system ***Kirkby et al***. (***2013***). In sensory systems, topographic relationships initially established by chemotrophic gradients are reinforced and refined by spontaneous activity in the periphery, which provides positional information because synchrony among sensory inputs drops off with distance. In the visual system, this activity takes the form of spontaneous retinal waves that propagate in 2D space and “synchronize” the firing of nearby neurons, driving visuotopic refinement in the thalamus, colliculus, and cortex (***Seabrook et al., 2017**; **Huberman et al., 2008***).

While the mechanisms of correlated activity generation are well studied, the central mechanisms by which this activity is processed and transformed into synaptic plasticity and, ultimately, circuit structure are poorly understood. One crucial question is how the correlation timescale influences developmental plasticity and how correlation timescales are regulated within early circuits, because timescale can determine the plasticity mechanisms potentially available to the developing neurons (***Zenke and Gerstner, 2017**; **Drew and Abbott, 2006***). In adults, thalamic and cortical neurons produce precise correlations on the timescale of milliseconds, which depend on the visual stimulus (***Butts et al., 2007b***). By contrast experimental and theoretical work has shown that during development information on visual topography during retinal waves is conveyed in coarse-grained correlation and vanishes in time windows below 100 ms (***Butts and Rokhsar, 2001**; **Butts et al., 2007a***). Correlation below such timescales is likely to be damaging as it has the potential to induce synaptic plasticity based on non-informative activation. Early synapses make several adaptations to accommodate the long timescales of development. These include synaptic currents which are slower to increase the integration window (***Hauser et al., 2014***), and burst-time-dependent plasticity with characteristic time-windows around 1 second to refine topographic relationships (***Butts et al., 2007a***).

How early synaptic properties contribute to synchronization has not been intensively studied. The promiscuous and imprecise synaptic connections observed during early development should drive correlation among target neurons, even if their activity is non-informative. For example, in the visual thalamus (dorsal lateral geniculate nucleus dLGN), each relay neuron receives functional inputs from 10 (***Jaubert-Miazza et al., 2005**; **Bickford et al., 2010***)] to 20 (***Chen and Regehr, 2000***) nearby retinal ganglion cells (rGC) on postnatal days 7-10 (P7-P10), prior to refining to the 1-3 seen in adults. Such polysynaptic convergence should cause correlation among relay neurons and their cortical targets with high temporal precision (***Sailamul et al., 2017***). Calcium imaging shows high correlations in the visual cortex during early development, which decorrelate rapidly around eye-opening (***Rochefort et al., 2009**; **Siegel et al., 2012***), but these correlations are present only at the timescales of retinal waves. Rapid timescale correlations are absent in the cortex and thalamus during early development, emerging only after eye-opening (***Colonnese et al., 2017***). The absence of precise spike correlation in dLGN activity at this age raises questions: Are there synaptic or circuit mechanisms that prevent relay neurons from synchronizing at timescales below the informative timescales of retinal waves? What functional advantage accrues by preventing the thalamus from the precise correlated activity?

Here, we address these questions using a biophysically detailed model of heterogeneous neurons in dLGN at P7-P10 driven by spike-trains of retinal ganglion cells recorded *ex-vivo* at these ages. We show that levels of convergence expected at these ages can cause high-levels of precise correlations among the modeled neurons at timescales below those that convey spatial information on retinal position during the retinal waves, i.e. causes informationless “parasitic” correlation. Such parasitic correlations are specifically suppressed when developmentally appropriate ratios of NMDA/AMPA receptor currents are introduced at the retinogeniculate synapse. Using the same model, we show that adult levels of the recurrent cortical and thalamic reticular inputs would also reinforce precise correlation in dLGN, given the extensive convergence of the retinal axons. However, levels of input consistent with the ages studied here do not correlate relay neurons suggesting that the delayed development of these inputs relative to the retina keeps them below levels required for synchronization of TC neurons. Applying information theory analysis, we show that spikes of the dLGN model with parasitic correlation lose spatial information, which should be detrimental for the refinement of thalamocortical connections. Finally, using spikes of the dLGN model as inputs to the state-of-art model of plastic, homeostatically regulated, balanced excitatory-inhibitory cortical network (***Wu et al., 2020***), we exemplify possible consequences of aberrant large and fast-neuronal correlations on for the development of cortical circuitry. Our results suggest a novel developmental principle: early synapses and networks eliminate correlations resulting from immature refinement that would damage network formation.

## Results

### Reproduction of the excitability and heterogeneity of thalamic relay neurons in a biophysical model

To model spike correlation in the developing thalamic network, it is necessary to accurately reproduce neuron dynamics, synaptic properties, and network heterogeneity at the corresponding age. These models must be heterogeneous to avoid synchronization strictly because neurons are identical; therefore, if spike correlation is observed, it would not be due to the homogeneity of neurons in the network. Even for relatively small networks, model heterogeneity requires up to a few hundred different neuron models to populate the network. Therefore, we used the standard approach of evolutionary multiobjective optimization (EMO) to constrain model dynamics to match the dynamics observed in our electrophysiological data (***Neymotin et al., 2017**; **Dura-Bernal et al., 2017***). EMOs allowed us to create a database of models that reproduce essential features of thalamocortical neurons at P7-P10.

The neuron membrane dynamics were derived from 10 thalamocortical (TC) neurons recorded at P7-10 *in vitro*, see Experimental Procedures section. Even adult TC neurons are relatively electrically compact (***Sherman and Guillery, 2004***; ***Bloomfield and Sherman, 1989***). Developing TC neurons have shorter and thicker processes (***Charalambakis et al., 2019**; **El-Danaf et al., 2015***), which allowed us to use a conductance-based “pen-and-ball” two-compartment model with a single segment for the somatodendritic compartment and a multisegment compartment for an axon. We adopted a state-of-art model of young adult (P14-18) somatosensory thalamocortical neurons in the ventrobasal thalamus developed by ***Iavarone et al***. (***2019***). We use state-of-the-art (genetic algorithms with nondominated sorting) and developed in-house (genetic algorithm with Krayzman’s adaptive weights) EMOs to fit dynamics of somatic voltage recorded in a current-clamp protocol. Both EMO methods yielded similar quality and quantity of acceptable models and therefore were used independently to avoid a potential bias of a single optimization method. Details of the second method are given in the appendix Genetic algorithm with Krayzman’s adaptive weights (KAMOGA).

Because neuron geometry, expression of specific subunits for ion channels, and densities of these channels change during development, in addition to the standard free parameter set for fitting, such as channel densities (conductance) in the somatodendritic compartment, we allowed minor adjustment in the additional classes of model parameters, such as:

- intracellular calcium buffer depth,
- calcium pump rates,
- reversal potentials for sodium, potassium, and nonselective voltage-gated cation channels (h-channel, HCN)
- time constant for the calcium-activated potassium channel (SK)
- half-points of Boltzmann’s steady-state functions
- geometry of both compartments – the length of the somatic compartment and the length and diameter of the axonal compartment

Thus, in total, the model had 29 free parameters for optimization, listed in the Neuron Optimization Pipeline section in methods and supplementary dataset for Figure 1. The list of objective functions for fitting and description of full pipeline fitting-validation-evaluation is also given in the same method section.

**Figure 1.**
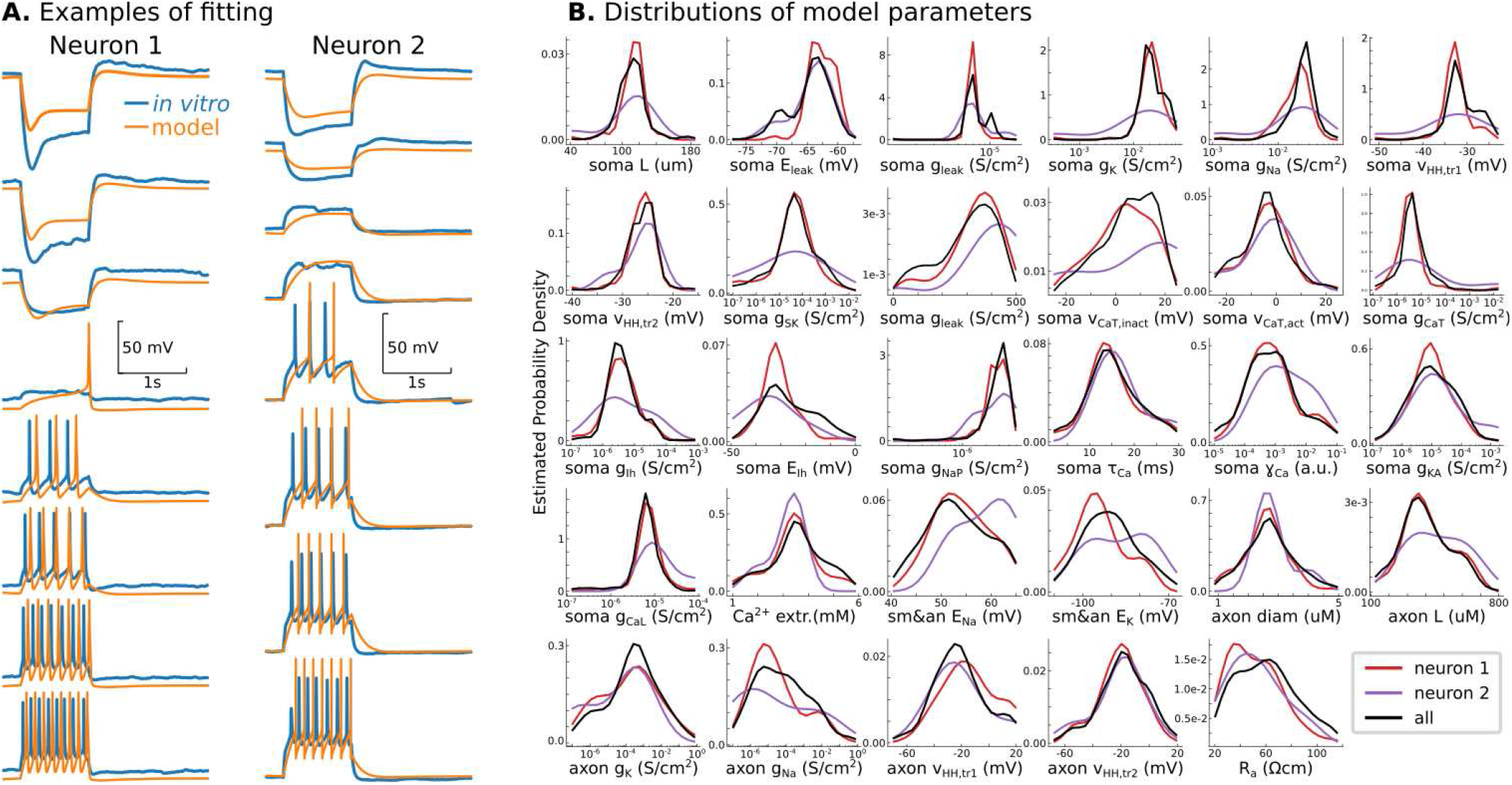
Conductance-based model of P7 TC neurons. **A** Randomly selected examples for two recorded neurons in the database. **B**. Probability density distributions of all 29 model parameters, obtained as Gaussian estimator with the bandwidth defined by Scott’s Rule. The kernel density estimator is implemented in the *scipy* Python library. Note that EMOs use a logarithmic scale for some parameters, and therefore, probability density distributions are also estimated in the logarithmic scale. Black lines are PDFs for all models in the database, and color lines indicate PDFs for models fitted to recorded neurons shown in A **Figure 1—figure supplement 0—source data 1.** Data base with 286 models of TC neurons at P7 https://github.com/rat-h/DevelopmentOfThalamocorticalNeurons **Figure 1—figure supplement 1.** Principal component analysis shows the separation of models fitted to the same recorded neurons as in Figure 1. The color code is the same as in Figure 1B in the main text. PCA decomposition was performed using *sklearn* Python library.

Both EMOs yield from 20 to 100 acceptable but different models for each recorded neuron, similar to the outcomes reported by ***Iavarone et al***. (***2019***) and ***Neymotin et al***. (***2017***). Therefore, these fitting procedures allow to accumulate a few thousand models in our current database. Figure 1A shows two models for two different recorded neurons, randomly chosen from the database. A reduced, fully validated, and human-evaluated database of 286 models, enough to reproduce any results in this paper, can be found in the supplementary data for Figure 1.

Somatic voltage dynamics in the current-clamp protocol may be insufficient to specify the unique ion-channel compositions of conductance-based models (***Marder, 2011**; **Prinz et al., 2004**, **2003***). Because multiple channel configurations can underlie similar model dynamics, it is expected that EMO, extensively searching for model configurations, converges to one of the regions where model dynamics match the target dynamics of the recorded neuron. For each recorded neuron, we ran from 2 to 20 independent EMOs, each starting from the random set of points in the parameter space. Therefore, we expected to find multimodal distributions of model parameters with peaks at convergence regions. Although the dynamics of TC neurons are different, parameters in the database do not show multimodal distributions (Figure 1B). Moreover, principal component analysis of parameters in the resulting database shows that, with a few exceptions, the models fitted to the different recorded neurons separate well in the principal component space (Figure 1 – Figure Supplement 1). Surprisingly, PCA indicates distinctive correlations of model parameters (PCA features) that characterize each recorded neuron, suggesting that EMOs achieve precision when recorded neurons can be distinguished by the parameters of their models.

Thus we conclude that our database is a valid representation of the real dLGN TC neurons at the specific point of their development. We expect the models have somewhat greater heterogeneity than real neurons due to the inability to determine a single exact parameter set for each recorded neuron.

### NMDA receptor dominance at developing retinothalamic synapses decorrelates relay neurons in millisecond timescale

We first examined the role of glutamatergic synaptic currents in the regulation of the correlation of thalamocortical neurons during the P7-10 period. As at many developing synapses, NMDA receptors provide the dominant conductance, even comprising the only receptor type in new synapses (***Rumpel et al., 1998**; **Chen and Regehr, 2000**; **Shah and Crair, 2008***). NMDA receptors are an established regulator of synchronization in adult networks (***Jacobsen et al., 2001***), and, thus, likely to play a critical role at these ages.

To systematically study the role of synaptic currents in precise spike correlation, we used our database to construct a small model of the dLGN network with realistic heterogeneity. This network is driven by rGC cell spikes obtained from the *waverepo* repository of early retinal activity recorded *ex-vivo* (***Eglen et al***. (***2014***), https://github.com/sje30/waverepo). We selected recordings of P6-P10 wild-type mice, recorded with a high spatial-resolution electrode array, and at least 30 adjacent active electrodes to provide sufficient spatial sampling and resolution ***Maccione et al***. (***2014***) and ***Stafford et al***. (***2009***). Recurrent inputs from the visual cortex provide an indispensable excitatory drive to dLGN at this age (***Murata and Colonnese, 2016***), and inhibitory inputs from the thalamic reticular nucleus (TRN) are also likely to be active (***Evrard and Ropert, 2009**; **Minlebaev et al., 2011***). These inputs were modeled as non-specific feedback in the sections below and ignored here for direct examination of the NMDA role. While present in the dLGN at this age, intrinsic interneurons are not reliably driven by retinal input before eye-opening (***Bickford et al., 2010***), and so we did not include them in the model.

To set the synaptic parameters for the retinogeniculate synapse, we used the ratio of peak-to-peak AMPA to NMDA currents *I_AMPA_/I_NMDA_* = 0.78 ± 0.09 according to the closest published measurement (P7) (***Shah and Crair, 2008***). This observation allows estimation of the ratio of peak-to-peak conductance for these currents at *g_NMDA_/g_AMPA_* ≈ 2.25 (see details in the dLGN Network Model method section). The retinogeniculate synapses also exhibit a strong paired-pulse ratio of 0.73 (***Chen and Regehr, 2000***), which we model using the simplified Tsodyks-Markram model (***Tsodyks et al., 2000***). The single RGC fiber synaptic conductance is unknown, though it appears to be less than required to fire a TC neuron (***Dilger et al., 2011**; **Liu and Chen, 2008***). Instead of setting the synaptic conductance into some specific value, we assumed that total synaptic conductance is under control of a homeostatic process with a target set-point of average firing rate and allowed homeostasis to regulate the firing rate at the level of individual neurons. For these simulations, the firing rate set-point was 0.5 spikes/s, which accounted for a mean firing rate in vivo of 1 spike/s (***Murata and Colonnese, 2018***), and the fact that blockade of the corticothalamic feedback reduces this rate by 50% (***Murata and Colonnese, 2016***).

During the first 3-4 weeks of development, the functional convergence of RGC axons on single relay neurons is reduced from >20 to 1-3 (***Guido, 2018**; **Liang and Chen, 2020***). The estimates of how many rGC axons converge to a single TC cell at P7-P10 vary from 10 (***Jaubert-Miazza et al., 2005**; **Bickford et al., 2010***) to 20 (***Chen and Regehr, 2000***). Because convergent inputs are most likely to come from adjacent RGC (***Liang and Chen, 2020***), we organize retinogeniculate synapses such that both the connection probability and synaptic conductance have Gaussian dependence on the distance between rGC and TC neurons, with the degree of convergence set by the parameter *σ*. We test a range of *σ* from 1, which corresponds to adult convergence (1-3 rGC connections per TC neuron), to 9 (20 rGC per TC neuron; *σ* = 4 generates 10). For each *σ*, ten network models are generated for analysis. Each simulation was run until the total glutamatergic conductance reached the steady-state region and the mean population firing rate reached the homeostatic set-point of 0.5 spike/second within 10% accuracy. We computed the spike correlation in each model from the last 20 min of network activity (when synaptic weights were stabilized). Correlations were computed as in previous experimental studies (***Colonnese et al., 2017***): the spike train of each neuron is convolved with a Mexican-hat-like kernel (a difference between 20 ms Gaussian and 80 ms Gaussian), and the Pearson correlation coefficient is computed for each pair of neurons within the population.

When NMDA/AMPA ratios are set to biologically accurate levels, the distribution (Fig A2) and mean (Fig A3) of spike correlations are in good agreement with experimental observations for these ages (***Colonnese et al., 2017***). Mean correlations are around 0.02-0.03 for expected levels of convergence *σ*=4-9 at these ages. The distribution of correlation was centered near zero, and some pairs displayed positive correlation, as observed in the thalamus. While biologically accurate, the level of correlation is unexpectedly low given the levels of input convergence and showed a surprising insensitivity to the convergence parameter *σ*.

We hypothesized that this low correlation and convergence-insensitivity might be a result of the NMDA receptor dominance at immature synapses. To test this, we examined a network with only fast AMPA receptors at the rGC synapse. In this case, we observe a dramatic increase in mean spike correlations, which reached 0.3-0.4 (Figure 2B) at biologically relevant levels of *σ*. Furthermore, the convergence factor *σ* strongly controls spike correlation in these networks (Figure 2B2, B3). Moreover, the mean spike correlation is relatively low for the adult convergence (*σ* = 1) and shows narrow distribution with a peak slightly shifted from zero.

**Figure 2.**
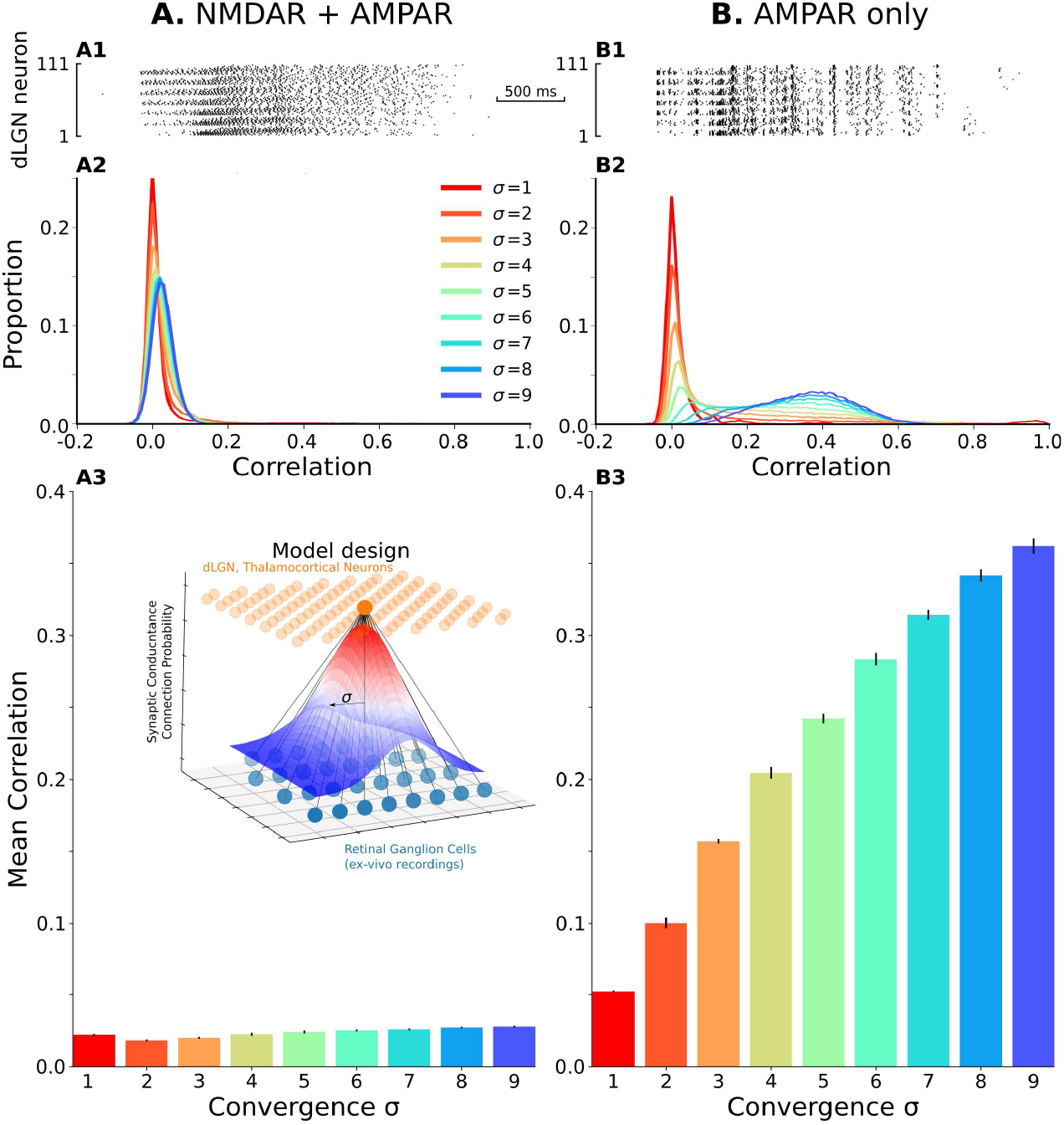
The model of the dLGN network at P7-P10 activated by spikes of rGC recorded *ex-vivo*. **A**. Retinogeniculate synapses have both NMDA and AMPA currents, with NMDA dominance. **B.** The same models with NMDA currents disabled. **A1, B1** examples of two model (*σ* = 4) responses to the same retinal waves in rGC firing. **A2 and B2** Distributions of pairwise spike correlation for different convergence factors (*σ*). **A3 and B3** mean and standard deviation of network-wise average pairwise spike correlation for ten network models for each type of the synaptic drive and each of convergence factor (*σ*). **The insert** schematically shows the model design.

These results show that slow NMDA receptor currents can desynchronize neural firing in a pure feedforward network, a phenomenon that has been well established for recurrent networks by ***Pinsky and Rinzel*** (***1994***), and may explain why the thalamic and cortical activity is so poorly correlated *in vivo* at rapid timescales at these ages (***Colonnese et al., 2017***), despite massive co-activation by the poorly refined retinal inputs (***Jaubert-Miazza et al., 2005**; **Bickford et al., 2010**; **Chen and Regehr, 2000***). In total, our model suggests that one of the critical roles of NMDA receptors is to prevent precise temporal correlation in the dLGN network, which otherwise would appear as parasitic correlation due to incomplete retinotopic refinement at P7-P10. As shown below, such synchronization could be detrimental because it loses relevant topographic information contained in retinal waves ***Butts and Rokhsar*** (***2001***), needed to refine thalamocortical projections. It also adds a correlation that does not represent any structure and can damage the upstream networks that will operate with the same time precision in adults (***Butts et al., 2007b***).

### Role of the thalamic reticular nucleus in precise spike correlation in dLGN

One objection to our hypothesis that NMDA receptors are abolishing the rapid correlations in the network because they would be deleterious to development is that the prevalence of NMDA receptors in retinogeniculate synapses is needed for synaptic plasticity and/or amplification (***Connelly et al., 2016***; ***Chen et al., 2002***; ***Liu and Chen, 2008***) and decorrelation is just an unfortunate “side effect” of these functions. We considered whether circuit mechanisms might exist to mitigate such a side effect and improve spike correlation in the developing dLGN.

One possibility is the recurrent inhibitory projections from TRN (***Pinault, 2004***). These GABAergic inputs are present and inhibit TC neurons by P7-P10 (silencing of TRN neurons at this age increases dLGN neuron firing by 2.3 fold (MTC, unpublished data)). Unfortunately, to our knowledge, direct measurements of TRN→dLGN synaptic conductances and delays do not exist for these ages. TC neurons do not establish synaptic connections to TRN neurons until the late second postnatal week, even though they pass TRN on the way to the cortex (WG unpublished data), suggesting that recurrent TRN input must arrive through the corticothalamic loop at a long delay (Figure 3A). To maximize the potential effect of TRN inputs on precise spike correlation, we modeled them as non-specific (to all neurons) inhibitory synapses. We then systematically studied synaptic conductance and delay parameter space to gauge the full potential of this projection to influence dLGN activity. Again, for each conductance-delay pair, ten different models with the same biological NMDA/AMPA conductance ratio of 2.25 were created, and each model ran until the mean firing rate of the population reached the homeostatic set-point 0.5 Hz ± 10 (Figure 3). We study an exhaustive range of conductance-delay pairs, many of which are likely far from biologically plausible values. To determine which pairs result in firing changes to dLGN neurons similar to that observed in vivo when TRN is silenced, we disable both the homeostasis and TRN feedback in models that reached homeostatic steady-state and measured firing rates to find the contour line where this ratio matches the experimental value *F_TRN–_/F_Control_* = 2.3. These contour lines overlapped on correlation heatmaps are shown in Figure 3B and C.

**Figure 3.**
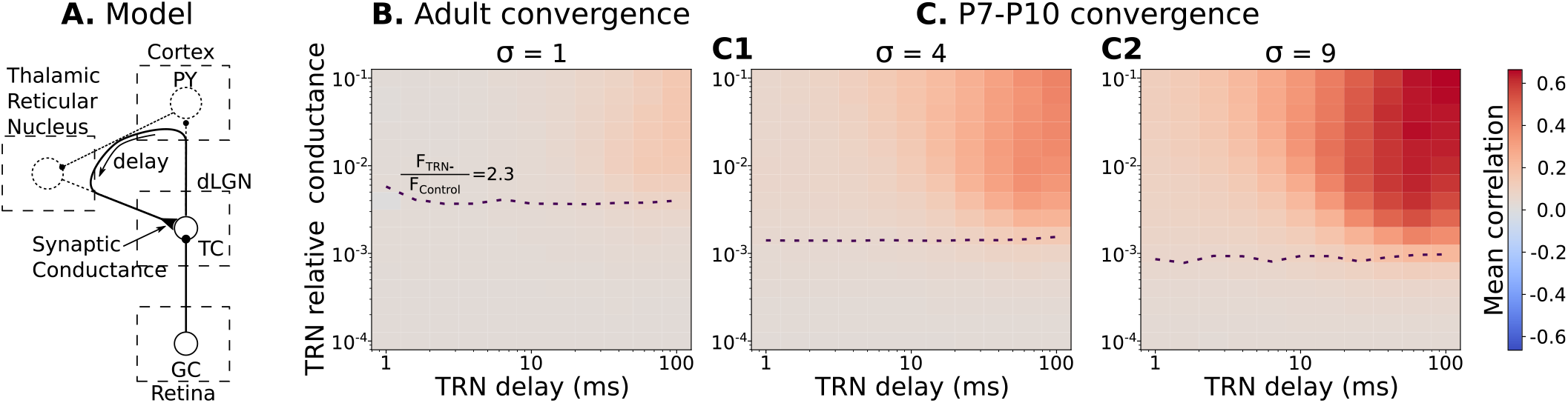
Parameters of TRN inhibitory feedback, that match *in vivo* observations, are outside the range when they can induce precise correlation in TC neurons. **A.** Schematic of the model design. **B.** Effect of different levels of TRN relative inhibitory conductance (ordinate) and delay (abscissa) on mean spike correlations in a model with adult-like convergence of rGC inputs to a single TC neuron (*σ* = 1). Estimated effect of TRN silencing on firing rate is shown as a dashed line. **C.** The same as in B but for two estimates of P7-P10 convergence: 10 inputs per TC neuron (*σ* = 4, C1) and 20 inputs per TC neuron (*σ* = 9, C2).

For adult convergence *σ* = 1, inhibitory TRN connections do not drive significant correlation under any sampled conditions. Therefore, this feedback is safe and cannot corrupt information conveyed in the fine-grain timescale in adults (***Butts et al., 2007b***). Even for strong and significantly delayed inhibitory currents, the mean spike correlation is below 0.1 (Figure 3B). By contrast, at P7-P10 convergence levels (*σ* = 4 or *σ* = 9), some combinations of high-conductance/long-delay (>10ms) can increase mean spike correlations (0.6 and higher) (Figure 3C1, C2). However, none of the parameter pairs that resulted in large correlations matches an increase in the firing rate *F_TRN–_/F_Control_* = 2.3 observed *in vivo*. This modeling suggests that the delayed development of functional TRN synapses, as observed (***Murata and Colonnese, 2016***), is desirable in part because strong TRN synapses early in development (particularly slow ones) could re-introduce dLGN correlations that result from the early poly-innervation.

### Role of the combined cortical and thalamic reticular nucleus feedback on precise spike correlation in developing dLGN

Although realistic levels of TRN inhibitory synaptic conductance are not sufficient to induce high spike correlations in the dLGN networks, there is still the possibility that the excitatory cortical feedback (possibly in combination with inhibitory inputs from TRN), can increase spike correlation of TC neurons to compensate for the NMDA receptor induced desynchronization. To test whether a combination of excitation and inhibition can correlate TC neuron spiking, we add an excitatory feedback loop that represents the cortical effect on dLGN (Figure 4A).

**Figure 4.**
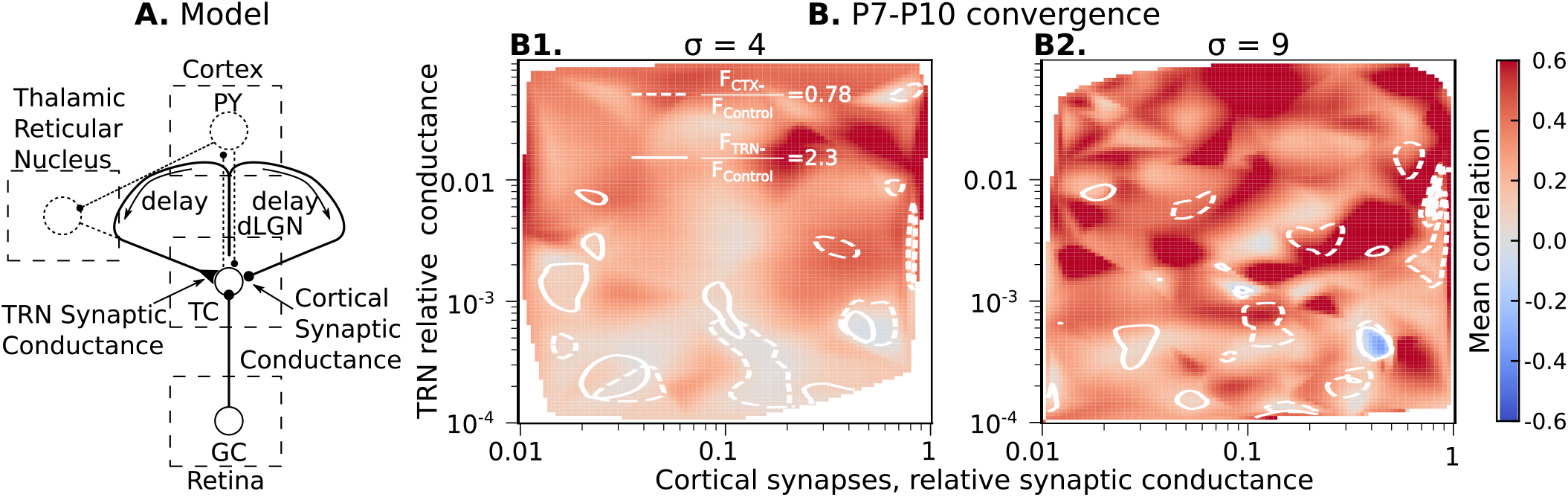
Parameters of cortical excitation and TRN inhibition, that match *in vivo* observations, are outside values where they can synchronize TC neurons. **A.** Schematic of the extended model. **B.** Dependence of mean network correlation upon TRN and cortical relative synaptic conductance. Two heatmaps with the convergence of 10 inputs per TC neuron (*σ* = 4, B1) and 20 inputs per TC neuron (*σ* = 9, B2) are shown. White solid lines and white dash lines indicate a mean change in firing rate which match observed in TRN silencing (*F_TRN–_*/*F_Control_* = 2.3) and in the cortex silencing (*F_CTX–_/F_Control_* = 0.78) *in vivo* experiments, correspondingly. These show the regions of possible conductance based on *in vivo* values. **Figure 4—figure supplement 1.** Heatmaps for mean spike correlation (left set), *F_TRN–_/F_Control_* (middle set), and *F_CTX–_*/*F_Control_* (right set) for all 4 model parameters: TRN synaptic conductance *g_TRN_*, TRN delay, CTX synaptic conductance *g_CTX_*, and CTX delay. Convergence *σ* = 4. **Figure 4—figure supplement 2.** The same as in Figure 4–Figure Supplement 1, but for *σ* = 9

As for TRN, exact measurements of CT feedback conductance and delay do not exist. Therefore, we evaluate the roles of cortical synaptic conductance (*g_CTX_*), cortical feedback delay, TRN synaptic conductance (*g_TRN_*), and TRN feedback delay to dLGN synchronization by treating each as free model parameters. For each parameter set, ten models were generated as above, and simulation for each network model runs until the mean firing rate of the population reaches the homeostatic set-point. In this case, the set-point was 1 spike/s, the level observed in vivo because the CT feedback was incorporated. Homeostatic regulation has access to both excitatory inputs, therefore, represents heterosynaptic plasticity (***Fiete et al., 2010**; **Wu et al., 2020***).

With four independent axes, the model parameter space is too big to explore systematically on available computational resources. Instead, we used Monte-Carlo sampling and reconstructed maps of mean spike correlation, mean change in firing rate when TRN is silenced *F_TRN–_*/*F_Control_*, and mean change in firing rate when the cortex is silenced *F_CTX–_/F_Control_* (Figure 4 supplementary 1 and 2). Although the parameter space is four-dimensional, we found two critical parameters to primarily define necessary conditions for TC spike correlation: conductance of TRN and CTX synapses. We project all sampling points points onto the two-dimensional conductance map to simplify the visual representation. To visual regions where the models generated CT and TRN inputs, that when blocked have similar amplitude effects on firing, we overlap contour lines for *F_TRN–_/F_Control_* = 2.3 and *F_CTX–_/F_Control_* = 0.78 on mean spike correlation. This analysis shows that while there are parameters or CT and TRN inputs that generate high correlation even in the presence of NMDA receptor dominance, these invariably lay outside the intersections of the two contour lines. Therefore, combinations of synaptic conductance that reproduce the *in vivo* data are insufficient for network synchronization. These data support our hypothesis that the developing thalamocortical system avoids configurations that would drive precise correlation resulting from the exuberant retinal inputs.

### Precisely correlated dLGN spikes lose retinotopic information

To see whether precisely correlated spikes of thalamocortical neurons convey spatial information and can be used for the refinement of thalamocortical connections, we applied classical information theory analysis suggested by ***Butts and Rokhsar*** (***2001***). The essence of this approach is to measure mutual information between interspike/interburst intervals (ISI, Δ*t*) for spikes of two neurons and distance (*r*) between these two neurons (*I*[*r*, Δ*t*]) which quantifies information about distances conveyed by neuron spiking (Figure 5A, see method section Quantification of mutual information). We computed mutual information *I*[*r*, Δ*t*] for dLGN models with different convergence factors (*σ*) described in detail in section NMDA receptor dominance at developing retinothalamic synapses decorrelates Figure 5B and C shows the effect of synaptic current composition on spatial information in spiking of dLGN neurons. For adult convergence (*σ* = 1) spikes of neurons driven by pure AMPAR currents convey more spatial information than with the mixture of NMDA and AMPA receptors found in developing dLGN. In contrast, for both estimates of of P7-P10 convergence: 10 inputs per TC neuron (Figure 5 C1) and 20 inputs per TC neuron (Figure 5 C2) spatial information is much higher for the biological mixture of synaptic currents than with AMPARs only.

**Figure 5.**
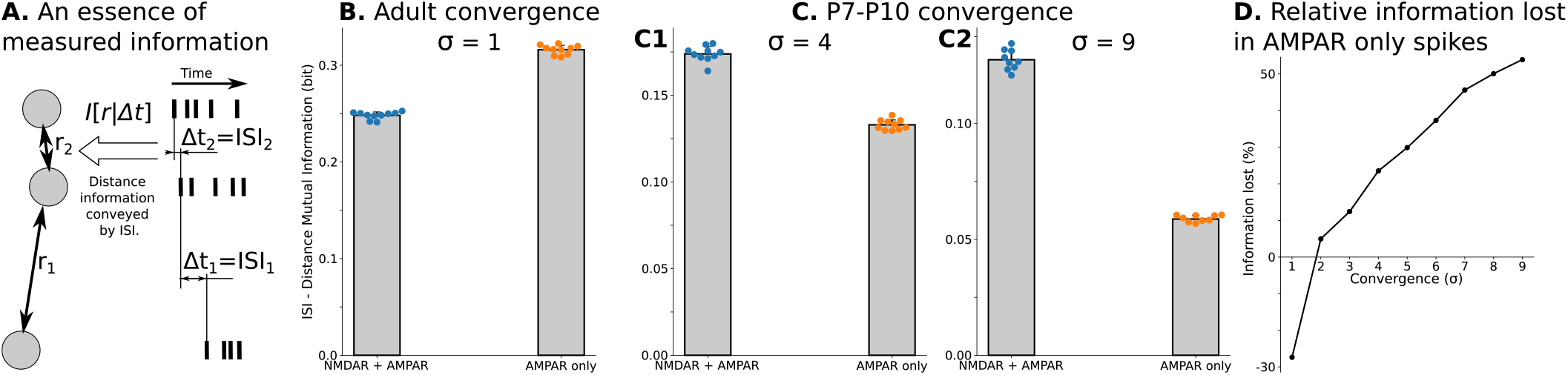
Spatial information encoded in interspike/interburst intervals (ISI) of TC neurons in the dLGN model. **A.** Mutual information *I*[*r*, Δ*t*] as a quantitative measure for predictability of distances *r* from observed ISI Δ*t*. **B.** Mutual information *I*[*r*, Δ*t*] in a model with adult-like convergence (*σ* = 1). **C.** The same as in B but for two estimates of P7-P10 convergence: 10 inputs per TC neuron (*σ* = 4, C1) and 20 inputs per TC neuron (*σ* = 9, C2). **D.** Dependence of information lost/gained in spikes of models with AMPA receptors only compared to models with NMDA+AMPA mixture of receptors on the convergence of rGC inputs to a single TC neuron (*σ*).

We qualify information loss or gain with AMPAR-only synapses as follows:

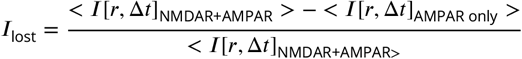

where < > denotes an averaging over 10 trials. This analysis (Figure 5D) shows that AMPAR-only synapses are advantageous only for adult convergence. For P7-P10 convergence, models with AMPAR-only synaptic currents lose from 25% to more than 50% of spatial information present with the natural composition of MNDAR and AMPAR currents. Overall, these results clearly indicate that the presence of parasitic correlations in P7-P10 (i.e., when it is not eliminated by NMDARs) reduces spatial information in TC neuron spikes and, therefore, can have a detrimental effect on both the speed and precision of thalamocortical projection refinement.

### Fast correlations, resulting from early input convergence, could damage cortical circuit formation

Finally, we wanted to ask how the NMDA receptor desynchronization of thalamic activity might help cortical development. We hypothesize that the parasitic correlations eliminated by the slow NMDA receptor currents are biologically disadvantageous for the formation of intracortical connections too. The parasitic correlation arises from the ratio of numbers of retinal neurons to TC neurons and the convergence of multiple rGCs onto a single thalamic neuron, the second condition do not exist in the adult. Because the specialized response properties such a direction selectivity and on/off identity have not emerged yet (***Wong, 1999***), and the critical topographic information represented in the retinal waves is conveyed only above 100ms (***Butts et al., 1999***), any plasticity driven by rapid early correlations is likely to not reflect the desired adult network connectivity. However, recent theoretical and computational research shows that correlated inputs from a subset of Poisson inputs to plastic and homeostatically regulated network can form a neuronal ensemble (a cluster) in an otherwise homogeneous network by potentiating and depressing intranetwork connections (***Gjorgjieva et al., 2011**; **Litwin-Kumar and Doiron, 2014***). Moreover, neurons in these ensembles show correlated activity, too (***Gjorgjieva et al., 2011***). Therefore, millisecond correlation in TC spikes should affect connectivity and activity in cortical networks, but this plasticity would be based on parasitic correlation, not spatial information.

To explore this question, we used a state-of-art model of the young adult visual cortex, widely used to model essential mechanisms of plasticity and homeostasis, clustering, and correlated activity in monocular deprivation experiments during the P24-P31 critical period (***Wu et al., 2020***). Although the dynamics of cortical activity during the earlier developmental stage, we examine here, are different from those in the age for which the model is optimized (***Colonnese, 2014***; ***Colonnese and Phillips, 2018***), it is sufficient to demonstrate how the different levels of correlation, we found, effects a plastic, homeostatically regulated network. Previous works examined the model with inhibitory-excitatory anisotropy by exclusively connecting subgroups of excitatory neurons with a reciprocal subgroup of inhibitory ones (***Wu et al., 2020***) or by strengthening excitatory-excitatory connections within a cluster (***Litwin-Kumar and Doiron, 2012***). However, our primary focus was on the group formation in isotropic networks. Therefore, here 800 excitatory and 200 inhibitory neurons are randomly distributed on a unit square, and the probabilities of connections between any pair of excitatory-inhibitory, inhibitory-excitatory, or inhibitory-inhibitory neurons were given a Gaussian dependence on distance. The parameter for distance dependence (*σ*) was fit to provide the same 20% connection probability as in the original model (***Wu et al., 2020***). Excitatory-excitatory neurons are connected randomly, with the same connection probability at 20%. To avoid edge-effect, the boundaries of the unit square are cyclical, imitating the continuation of the cortex in all directions.

Several parameters of the original model were adjusted for the earlier developmental stage P7-P10. Both resetting and resting potentials were set to −62 mV, as observed experimentally (***Colonnese, 2014***). Furthermore, the intracellular chloride concentration in immature neurons increases the reversal potential for GABAergic synapses, providing slight excitation or at least shunting inhibition (***Kirmse and Zhang, 2022**; **Murata and Colonnese, 2019***). Therefore, we set the reversal potential of inhibitory synapses to *E_inh_* = −62 mV. We added a magnesium voltage dependence for the NMDA receptor synaptic conductance to imitate the NMDA receptor magnesium block. Finally, because at P7-P10, the visual cortex is almost entirely silent if the thalamus is blocked (***Murata and Colonnese, 2016, 2018***), we removed any sources of input spikes, except those derived from the thalamic model.

We drive the same cortical model by spikes obtained from the thalamic model with and without NMDA receptor synaptic currents. As described in the NMDA receptor dominance at developing retinothalamic section above, that corresponds to TC neurons firing with low and high spike correlation, respectively. The simulations were run for 2.7 hours of model time, enough for synaptic plasticity, synaptic scaling, and homeostatic regulation to reach the steady-state. Note that we intentionally do not disable NMDA receptor synaptic current in the cortical model even when driven by spikes from the dLGN model without NMDA receptors as we wanted to test specifically the effects of high correlation on the cortex, not the effect of NMDA receptor blockade *per se*. Moreover, this modeling might have an experimental approach for validating of the model predictions in NR1 (*Grin1*)-null mice (***Arakawa et al., 2014***).

Driving the cortical model with either level of precise correlation resulted in distributions for total *input* synaptic conductance for E → E connections that were nearly identical (see Figure 6-Supporting Figure 1). By contrast, the *outgoing* connections within the cortex were strongly affected (Figure 6A). In networks driven by correlated dLGN spikes (NMDA receptor blocked in dLGN), the distribution of total outgoing synaptic conductance for single neurons was stretched to both sides (Figure 6A). In the high-correlated dLGN input condition, up to a quarter of neurons did not have postsynaptic partners (total postsynaptic conductance became equal to zero, see Figure 6B1). In contrast, in networks driven by uncorrelated dLGN spikes (NMDA receptor active in dLGN) only a small fraction (7-10%) of such ineffective (“inconsequential”) neurons (Figure 6A left panel) were generated (Figure 6B1). On the other side of the distribution, driving the cortical model with highly correlated inputs resulted in 1-13 neurons with maximal conductance on almost all their postsynaptic partners (Figure 6B2). In the low-synchronization condition, only 3 neurons out of 9 models were in the same range, with only one neuron in each network. The emergence of multiple highly “influential” neurons is likely to result in a single dominant group of neurons that controls neural activity, increases mean correlation (Figure 6B3), and would reduce response diversity.

**Figure 6.**
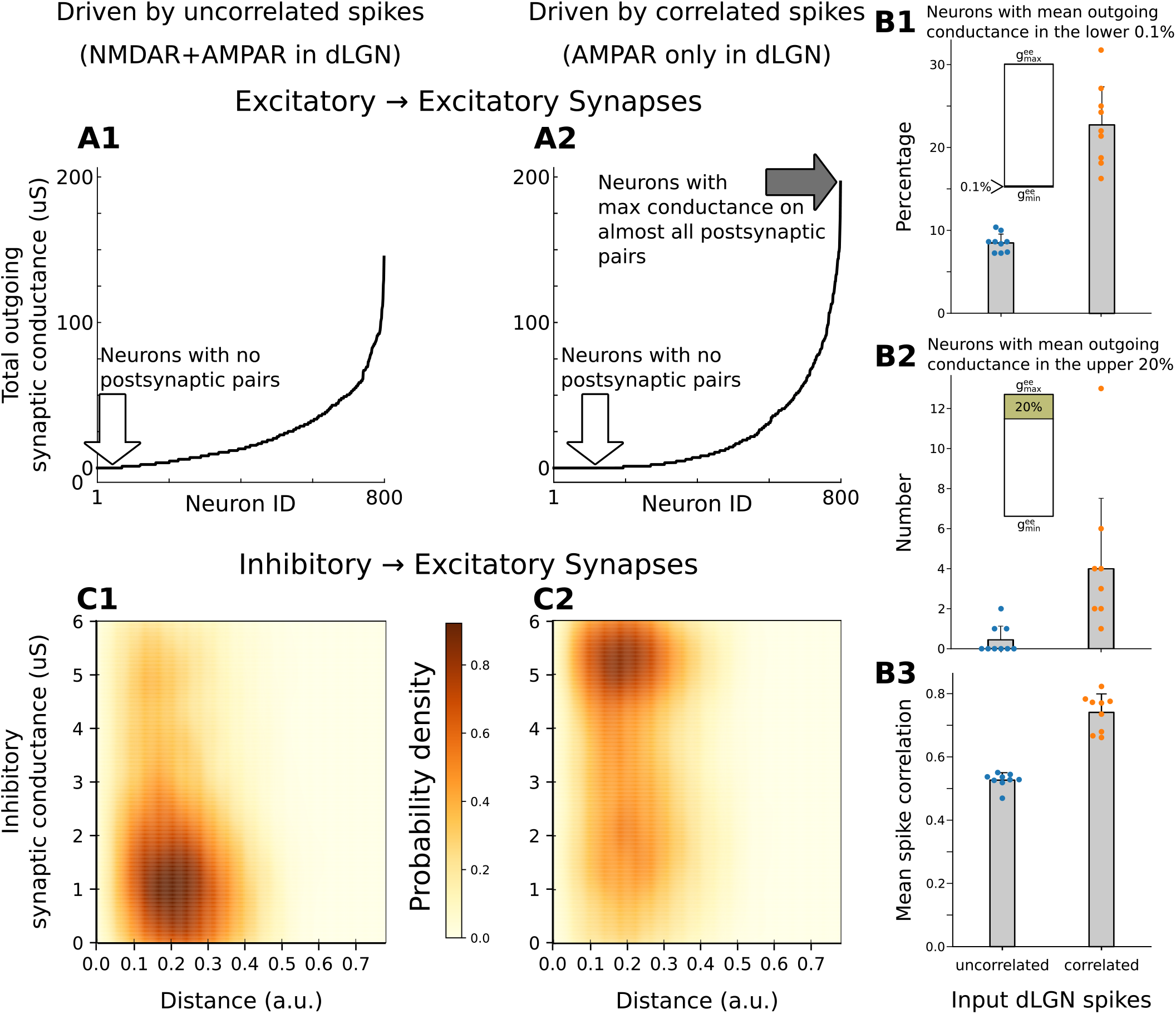
The difference in the connectivity and activity of plastic and homeostatically controlled networks activated by uncorrelated and correlated dLGN spikes. **A.** Distributions of total synaptic conductance on postsynaptic neurons for a given presynaptic neuron. Two examples are shown for networks driven by the uncorrelated spike (A1) and correlated spikes (A2). Blank arrows show groups that lost connections to postsynaptic neurons. The filled arrow indicates a group of 3 neurons with maximal conductance on almost all postsynaptic neurons. **B.**: percentage of the excitatory neurons with mean outgoing synaptic conductance is in the lower 0.1% of the allowed range (B1), number of neurons with mean outgoing synaptic conductance is in the upper 20% (B2), and the mean spike correlation (B3). **C.** Distributions of inhibitory-excitatory synaptic conductance in the same models, shown as multivariable kernel-density-estimator with conductance and distance as independent variables. **Figure 6—figure supplement 1.** Distribution of total *input* conductance sorted from bottom up (on the left) along with matrix of excitatory-excitatory connections shown as heatmap for the same networks shown on Figure 6 A and B of the main text. **Figure 6—figure supplement 2.** Probability I→E conducntace for 8 models driven by uncorrelated (left) and correlated (right) dLGN spikes.

Interestingly, correlation in input activity also dramatically changes inhibitory connections. Both E→I and I→I connections are not plastic in these models. However, I→E inhibitory synapses are controlled by iSTDP plasticity (***Vogels et al., 2011**; **Litwin-Kumar and Doiron, 2014**; **Wu et al., 2020***). The probity density distributions of *g_EI_* in the distance-conductance space (Figure 6C) revealed a profound increase in inhibitory synaptic strength when correlated inputs drove the model. When driven by uncorrelated spikes, moderate inhibitory synaptic conductance 0.25-1.5 uS is most probable. However, models driven by the highly correlated spikes show the highest probabilities for inhibitory conductance around the maximal possible value of 6 uS. The two levels of input correlation did not change the distance at which synaptic strengths converged into a specific preferable value, which increases probability density in the middle range distances (0.1 −0.3 a.u.) for both. Univariate kernel density estimators for distributions of synaptic conductance (or just simple histograms) are skewed towards the strongest possible inhibitory synapses for models driven by correlated spikes, while models driven by uncorrelated spikes have single-modal distribution with a peak around 1 uS (see Figure 6-Supporting Figure 2). These effects were consistent for all runs of the model.

Overall, our modeling shows that the presence of precise, millisecond correlation in thalamic spikes in early development can create a single dominant group of neurons that control the rest of the population activity, disconnect up to a quarter of the population from their postsynaptic recipients, and impose strong inhibitory control that enforces firing in synchrony with the dominant group.

## Discussion

In this study, we modeled some of the key synaptic and circuit factors that regulate the precision of spike correlations in the developing dLGN. By employing state-of-art and novel methods of multiobjective evolutionary optimization, we reconstructed the dynamics of thalamocortical neurons in dLGN at postnatal days 7-10. We constructed a heterogeneous network composed of these realistic cellular models which represents the dLGN network at a specific developmental time point during initial circuit formation. We drove this network with spikes of retinal ganglion cells recorded *ex-vivo* at similar ages to identify a previously unexpected role for NMDARs during development. Beyond their expected role in synaptic plasticity and amplification of weak synapses, we showed that NMDARs actively prevent the correlation of thalamic relay neurons on the milliscond-scale that would otherwise be generated by the extensive convergence of the refining retinal ganglion cells. We refer to these precise correlations as “parasitic” because they contain no information themselves and actually reduce topographic information conveyed by thalamic neurons to cortex. We further showed that if adult-like connectivity from the feedback projections of cortex and TRN were present, they would reinstate parasitic correlations, again given the extensive convergence of the retinal axons. However, at connection strengths observed at early ages, their effect is negligible. These results allowed us to hypothesize the desynchronization of thalamic neurons by NMDARs is an important component of early activity and that the developmental delays in cortical and TRN connectivity are likely selected by evolution to avoid the generation of precise, rapid-timescale, parasitic spike correlations in TC neurons. In addition to showing that precise neural correlations reduce topographic information, we test their effects on network formation using a plastic, homeostatically-regulated, cortical network model. This model showed that precise millisecond correlations in dLGN inputs reduces network diversity disconnecting of up to a quarter of the population from their postsynaptic recipients, increases the connectivity a small group of neurons and by drives strong inhibitory control of the network. Together these results suggest a detrimental loss of diversity of neural function and domination of the network by a small group of neurons. Overall, our results suggested a novel developmental principle that just as appropriate synchronization must be generated in inputs and their targets to refine synaptic connectivity (***Thompson et al., 2017**; **Feller, 2009***), parasitic synchronization arising “accidentally” from the exuberant connectivity present during early development must be minimized to maximize information transfer and prevent premature or aberrant plasticity in the downstream circuits.

### Why can precise correlation be detrimental? Dual effects of correlation during development

In the developing visual system, spontaneous waves of activation correlate activity of nearby ganglion cells (***Wong et al., 1993***). The spatial information supplied by this correlation is used to reduce exuberant connectivity and refine topographic mapping in the primary retinal targets as well as the visual cortex (***Huberman et al., 2008***). Genetic manipulation that increases the size and intensity of retinal correlation reduces the refinement of retinal projection in the thalamus and superior colliculus (***Seabrook et al., 2017***). Because waves propagate slowly and only poorly correlate the firing of rGCs, topographic information is conveyed by the developing retina only at timescales greater than 100 ms (***Butts et al., 1999, 2007a**; **Butts and Kanold, 2010***). Within the area of cortex driven by a single wave neurons maintain a diversity of neural receptive fields, including multiple orientation selectivity, direction selectivity, and even visual responsiveness, which is critical for visual processing. These diverse receptive fields emerge from refined thalamic as well as recurrent local cortical inputs (***Niell and Scanziani, 2021***). Our results suggest that active decorrelation of thalamic responses by NMDARs (which is supported by the delayed development of feedback inhibition and excitation) may be critical to the maintenance of this diversity. By eliminating correlations on timescales below those that convey relevant information at a particular age and by preserving the correlations in coarse time scales - information that are informative, the sysaptic and circuit mechanisms we identify allow for topographic refinement and stabilization before the emergence of a diversity of receptive fields within an area of visual space. We did not explicitly model single neuron membrane dynamics at later stages of development. However, it is likely that with decreasing NMDAR dominance and increasing decay times (***Liu and Chen, 2008***), as well as the formation of recurrent connections (***Guido, 2018***), the ability to support fast synchronization increases as visual information that uses such rapid timescales develops after eye-opening (***Usrey and Reid, 1999**; **Butts et al., 2007b***). Our results suggest a novel mechanism by which critical periods can be regulated: the changing timescales of synaptic currents and feedback connections modifying the time and degree of correlation. For example, reduced corticothalamic input could decrease retinothalamic refinement (***Hooks and Chen, 2020***) by reducing fast, local synchronization or inducing synchronization among nearby neurons.

### Multiple roles for NMDA receptors during development

An essential role for NMDARs in the segregation and refinement of glutamatergic afferents has been demonstrated in multiple species and sensory systems (***Ewald and Cline, 2009***). This role has largely been ascribed to their capacity to induce synaptic plasticity in response to synchronous activity. However, NMDARs are not critical for synapse stabilization, sprouting, or many forms of afferent refinement and segregation (***Iwasato et al., 2000**; **Hahm et al., 1991**; **Colonnese and Constantine-Paton, 2001**; **Huang and Pallas, 2001***), likely because other forms of calcium entry can drive the necessary synaptic plasticity for afferent refinement (***Lee et al., 2014***; ***Kuo and Dringenberg, 2012***). In addition to regulation of synaptic plasticity, NMDA receptors play important roles in network activity. NMDA currents amplify activity at the retinogeniculate synapse both through their long decay times (***Liu and Chen, 2008***) as well as their activation of plateau potentials (***Lo et al., 2002***) critical for afferent refinement (***Guido, 2018***). Here, we suggest a potential third role for NMDARs: the decorrelation of neurons on a fine-grain timescale while simultaneously enhancing slow correlations (***Mizuno et al., 2021***). We suggest that NMDAR dominance at the glutamatergic synapse is one of multiple adaptations to reduce synchronization, specifically fast-correlations, among neurons. While not specifically modeled here, heightened correlations among target neurons are expected following chronic NMDAR blockade. Such parasitic correlations are predicted specifically reduce the elimination and sorting of afferent axons that show some level of correlation (for example, same-eye ganglion cells) while not affecting those with very low correlation (for example, opposite eyes). Such an effect has been observed in the developing thalamus where NMDAR blockade prevents on-off segregation within the same eye, but not eye-specific lamination (***Hahm et al., 1991***).

For our modeling, we examined a model of developing LGN in which the loss of NMDARs was compensated for by homeostatic increases in AMPA receptor currents, to maintain a similar level of retinal drive. Blockade or knockout of NMDARs has been shown to increase spontaneous EPSCs (***Kesner et al., 2020***), increase AMPAR-driven circuit excitability (***Kesner et al., 2020***), and increase glutamatergic synapse density (***Rocha and Sur, 1995**; **Colonnese and Constantine-Paton, 2006***), consistent with our modeling. However, it is likely that *in vivo*, there are physiological limitations that prevent the developing nervous system from fully augmenting AMPAR currents in NR1 knock-outs to levels observed in our model. In fact, NMDAR knock-out or chronic blockade sometimes results in no change in AMPAR currents or expression (***Colonnese et al., 2003***) or their delay (***Zhu and Malinow, 2002***). Nevertheless, the insights provided here are still applicable regardless of the effects of NMDAR elimination of excitability because they demonstrate how NMDARs decorrelate early activity and provide testable predictions that can be calibrated to the levels of AMPAR homeostasis observed following particular experiments. We note that the decorrelation effect shown here are likely to be specific to circuits similar to the developmental age and configuration described here, specifically they require convergence from an input that synapses on many nearby neurons, is very sensitive to recurrent excitation and inhibition, as well as the slow membrane dynamics of the developing neurons.

### Model innovations and limitations

This study examined the mechanisms that abolish expected, but not observed, correlation in the developing dLGN. To this end, the dynamics of individual neurons, details of synaptic organization, specificity of retinal projections, and accurate representation of network heterogeneity are indispensable, requiring a detailed biophysical model of the developing thalamus, including reconstructing the dynamics of thalamocortical neurons at a specific point in the network developmental trajectory (P7). Simple current-clamp is insufficient for fully determining parameters for single compartment, single-segment, conductance-base models ***Marder*** (***2011***); ***Prinz et al***. (***2003***, 2004), and this limitation is greater for multi-compartment models, especially those including the axon compartment ***Almog and Korngreen*** (***2016***). Surprisingly, we found that almost all parameters show a unimodal distribution (Figure 1) and that models fitted to different recorded neurons could often be separated in principal-component space, suggesting that EMOs achieved accuracy sufficient to distinguish neurons by their model parameters (Figure 1—figure supplement 1). We found PCA of model parameters to be a useful tool for qualitative assessment of EMO quality and heterogeneity of obtained models.

The next step was to reconstruct the synaptic currents to a TC neuron. An extensive body of research allows the estimation of conductance ratio for NMDAR and AMPAR currents as well as the paired-pulse ratio for retinal inputs. To set the strength of retinal inputs we used the firing rate of TC neurons observed *in vivo* to “functionally” estimate synaptic conductance at this age by assuming that total synaptic conductance is subject to homeostatic regulation and firing rate is the target activity parameter (***Riyahi et al., 2021***). Using homeostatic regulation to set synaptic conductance is critical to model heterogeneous networks, where a single value for synaptic conductance would not work for all neurons. Although neurons have different mean firing rates even after homeostasis converges, these firing rates have plausible distribution around the mean firing rate. Therefore, by applying well-developed homeostatic mechanisms to a heterogeneous network, we acquired a novel approach that allows the maintenance of both the heterogeneity of the neurons and the heterogeneity of firing rates in the biologically reasonable ranges.

We reconstructed the dLGN network at the high detail level because, as shown in Figure 2, the interaction of synaptic currents and neuron intrinsic dynamics can dramatically change spike correlation in the dLGN network. However, there is no reason to model the cortical and TRN feedback in such detail. Although these inputs control firing rates of TC neurons, there is no evidence that they are precise, have well-established topographical alignment, or very reliable (***Murata and Colonnese, 2019***). Thus, we schematically modeled these feedbacks, introducing a novel hybrid detailed-phenomenological approach, where the synaptic and neuron levels are the detailed models, but TRN and cortical feedbacks are presented in a phenomenological fashion. Again using homeostasis and target firing rates to adjust and balance network excitability, we concluded that none of the feedbacks are strong enough for imposing millisecond spike correlation in dLGN networks.

To gain insight into why evolution suppresses parasitic correlation, we used a simplified cortical network model developed for older animals. It should be noted that neuron intrinsic dynamics at P7 and P24-P31 are very different. At early development, cortical neurons are slow, have broad action potential, and produce plateau-potentials in response to a retinal wave (***Colonnese, 2014***). Although we observed some kind of plateau-potentials in the leaky-integrate-and-fire model of cortical excitatory neurons, they appear due to a balance between excitation and inhibition which should not be present at this age (***Kirmse and Zhang, 2022**; **Murata and Colonnese, 2019***). Moreover, the spike correlation we report in this model (Figure 6C) is much higher than observed at this age *in vivo (**Colonnese et al., 2017***). Therefore, results obtained from the “cortical” model should be considered only as an illustration. For the further study of what kind of malfunctioning can be induced by activity with millisecond spike correlation, a model more closely resembling the cortical network at the specific point in the developmental trajectory is needed. However, this simple example shows that a network can undergo a significant transformation if parasitic correlation in TC activity is present, and such a transformation can be detrimental to the diversity of cortical ensembles.

### Novel principal of development

Our results unexpectedly suggest that the nervous system may have evolved to avoid specific network dynamics that appear due to the unrefined connections present during development. Many afferents refine their connectivity during development, a process that allows activity and experience to influence the final circuit connectivity (***Vonhoff and Keshishian, 2017**; **Kano and Hashimoto, 2009**; **Sanes and Lichtman, 1999***). How this exuberance affects the developing circuit dynamics and how circuits select which activity to use and which to discard has not been extensively considered. Our results suggest that just as specific circuit properties have evolved to generate and transmit activity necessary for proper circuit formation, such as retinal or cochlear waves (***Elstrott and Feller, 2010***), specific mechanisms, such as the dominance of NMDA current (Figure 2) or delay the development of TRN and cortical connections (Figures 3 and 4) developed to suppress activity that would be deleterious to the developing system. In this case, the detrimental activity is a parasitic precise correlation arising from early convergence in the unrefined inputs, but we expect there are other generators and other suppressors. Overall, we propose a potentially general principle of neurodevelopment: developing circuit and synaptic properties are balanced to optimize the transmission of informative activity as well as to suppress the dynamics which appear due to the incompleteness of network connections and which cannot be used to properly refine the network.

## Methods and Materials

### Experimental Procedures

*In-vitro* whole-cell patch recordings were obtained from dLGN neurons. Borosilicate glass pipettes were pulled from vertical puller (Narishige) and had a tip resistance of 5–10 MOhm when filled with an internal solution containing the following:117 mM K-gluconate, 13.0 mM KCl, 1 mM MgCl2, 0.07 mM CaCl2,0.1 mM EGTA, 10 mM HEPES, 2 mM Na-ATP, and 0.4 mM Na-GTP. The pH and osmolality of internal solution were adjusted to 7.3 and 290 mOsm, respectively. Brain slices were transferred to a recording chamber that was maintained at 35°C and continuously perfused with ACSF (3.0 ml/min). Neurons were visualized using an upright microscope (BX51W1, Olympus) equipped with differential interference contrast optics. Whole-cell recordings were obtained using a Multiclamp 700B amplifier (Molecular Devices), signals were sampled at 2.5–5 kHz, low-pass filtered at 10 kHz using a Digidata 1320 digitizer and stored on computer for subsequent analyses using pClamp software (Molecular Devices). Access resistance (15MOhm) was monitored continuously throughout the experiment, and neurons in which access resistance changed by 20% were discarded. A 10-mV junction potential was subtracted for all voltage recordings.

### Neuron Optimization Pipeline

Two methods of multiobjective evolutionary optimization (EMO) were used to reproduce the dynamics of TC neurons in the biophysical model. Namely: genetic algorithms with nondominated sorting (NSGA2) (***Deb et al., 2002***; ***Deb, 2001***), implemented in the *inspyred* Python library by Dr. Aaron Garrett (***Tonda (2020)***, https://github.com/aarongarrett/inspyred) and developed in-house genetic algorithm with Krayzman’s adaptive multiobjective optimization (see Genetic algorithm with Krayzman’s NSGA2 uses the Pareto archival strategy, selecting one model over another if it is better than or equal to the other model in all fitness functions and strictly better in at least one fitness function. This criterion is used to determine whether an individual model is selected for entry into the final archive, which only occurs if the model is at least as good as the other models in the archive.

NSGA2 performs well for single and multicompartment neuron models but requires pre-fitted passive cable properties, which is an additional step in the optimization procedure (***Neymotin et al., 2017***). In contrast, KAMOGA re-adjusts the weights of fitness functions so that the resulting distribution of overall fitness in the generation correlates with distributions of individual fitness functions (***Eremenko et al., 2019***). KAMOGA avoids over- or under-representing individual fitness functions in the overall fitness and balances EMO objectives during the optimization.

EMO ran 1024 generations of 240 models each (245760 models in total) either for NSGA2 or KAMOGA. The light elitist selection was used for KAMOGA, holding 30 best models in the next generation from the previous one (12%). Two sets of fitness functions were used: absolute difference in the number of spikes and Euclidian distance between voltages samples for a recorded neuron and a model. Both fitness functions were applied for selected traces in the current-clamp protocol, keeping the total number of fitness functions between 40 and 54, 20 to 27 traces, correspondingly.

After GA finishes, all obtained models are re-evaluated and sorted again. The top 510 models (0.2 %) are selected for automatic validation. Validation checks that model parameters are biophysically correct, namely: the length of an axon is longer than a diameter, the diameter of the axon is smaller than soma diameter, the depth of Ca^2+^ buffer is smaller than soma size, and that spikes propagate without decrement through the axon. Then the model sets that passes validation were evaluated by a human. The pipeline produces from 10s to a few hundred models for each recorded neuron.

The complete list of model parameters open for adjustment by EMO is given in the Open Model Parameters table. Note that EMO searches in logarithmic space for some model parameters, while the others have linear scale. The choice of scaling depends on the ratio between minimal(*p_min_*) and maximal(*p_max_*) boundaries for a particular parameter. For example, EMO performs better in a linearly scaled space for a reversal potential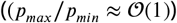, but shows better results in logarithmic scaled space for a channel conductance 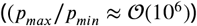.

The code for both EMO and supporting tools are available through the *GitHub* repository https://github.com/rat-

### dLGN Network Model

Parameter sets for individual TC neurons are randomly picked from the database. Each neuron model consists of a somatodendritic single-segment compartment and a multisegmental axonal compartment. The number of segments in the axonal compartment is set to an odd value so that segments are no longer than 0.1 of the AC length constant at 100 Hz (***Hines and Carnevale, 2001***). In the somatodendritic compartment, there are nine cross-membrane channels: leak current, fast sodium and delayed rectifier potassium currents, persistent sodium current, transient and depolarization-activated potassium current, L-type calcium current, low-threshold calcium current, SK-type calcium-activated potassium current, and nonselective voltage-gated cation current.

Neurons are deployed on 7×16 hexagonal lattice, 112 neurons in total. An ex-vivo recording electrode lattice (usually square) is scaled and center to the neuron lattice. Probability of connection between GCs and TC neurons is defined as exp(−|**r**_*GC*_, **r**_*TC*_ |^2^/*σ*^2^) while the synaptic conductance is defined as *g*_0_exp(−|**r**_*GC*_, **r**_*TC*_|^2^/*σ*^2^), where *σ* is a convergence parameter, *g*_0_ is the minimal synaptic conductance needed for triggering a single spike in neuron model for given parameter set and NMDA/AMPA conductance ratio, |**r**_*GC*_,**r**_*TC*_| is a distance between GC and TC neurons.

Each synapse is modeled as a two-stage process. The first is a simplified Tsodyks and Markram model (***Tsodyks et al., 2000***) implemented by Dr. Ted Carnevale (https://senselab.med.yale.edu/ModelDB/showmo The parameter of presynaptic single spike depression (*u*_0_) was set to 0.3 for match a paired-pulse ratio of 0.73 (***Chen and Regehr, 2000***). Time profiles for both NMDA and AMPA are modeled as double-exponential synapses with time constants 1 ms rise, 2.2 ms decay for AMPA (***Chen and Regehr, 2000***), and 1ms rise, 150 ms decay for NMDA (***Chen and Regehr, 2000**; **Dilger et al., 2015***). Proportion of NMDA and AMPA conductance was derived from the peak-to-peak ratio of currents *i_AMPA_/i_NMDA_* = *β_P7_* = 0.78 ± 0.09 (***Shah and Crair, 2008***). The current in for voltage clamp experiment is *i_vl_* = *g_vl_* * (*E* – *V_vl_*) where *E* = 0 reversal potential for NMDA and AMPA, and *v_VL_* potential for voltage clamp. Substitute the current and voltage to *g_vl_* = *i_vl_/v_vl_* we can get a fraction of AMPA to NMDA conductance as

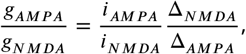

where Δ*_NMDA_* and Δ*_AMPA_* are differences between the holding and reversal potentials for NMDA and AMPA currents. For both ***Shah and Crair*** (***2008***) and ***Chen and Regehr*** (***2000***) experiments 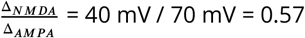, and ratio of NMDA to AMPA conductance, therefore

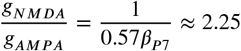

**Table 1.**
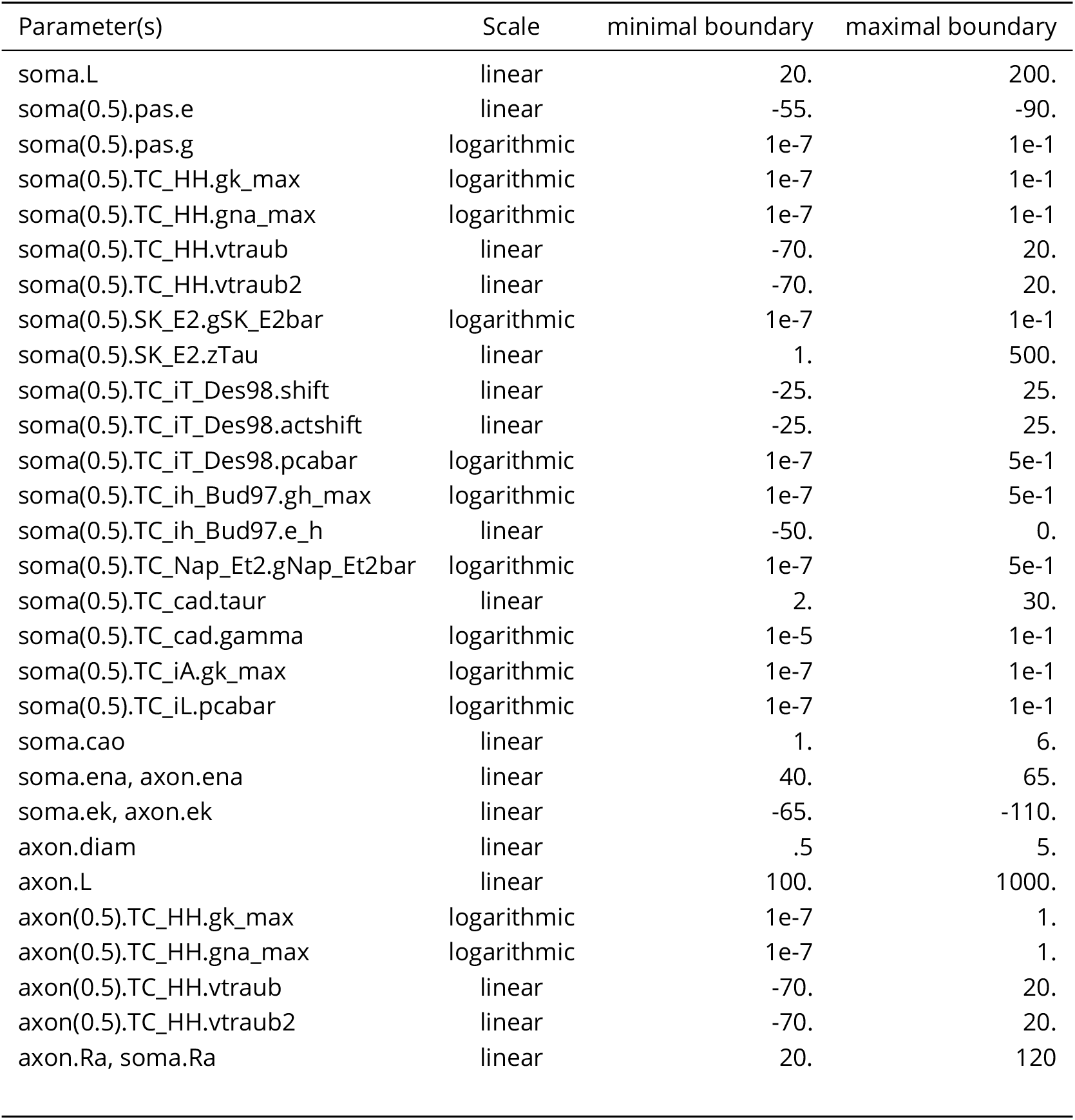
Open Model Parameters

Note that ***Dilger et al***. (***2015***) estimated *β_P10_* ≈ 1 which give *g_NMDA_/g_AMPA_*=1/0.57≈ 1.75, while the same value from ***Chen and Regehr*** (***2000***) *β_P10_* = 0.5 ± 0.1 gives *g_NMDA_*/*g_AMPA_*=1/(0.57 0.5)≈ 3.5. In this model, we used *g_NMDA_/g_AMPA_* = 2.25 value. We assume that 13% of NMDA current is conducted by Ca^2+^ ions and, therefore, we add this current as calcium current computed as Goldman–Hodgkin–Katz equation.

Because neurons in the dLGN model can show a wide range of firing rates, specifically just after model initiation, linear firing-rate homeostasis does not show robust results. We used nonlinear firing-rate homeostasis, which scales all synapses at the given neuron as

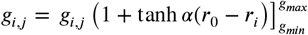

where *g_i,j_* is synaptic conductance from *j^th^* rGC neuron to *i^th^* TC neuron limited by maximal *g_max_* = 10*g*_0_ and minimal *g_min_* = 0 boundaries, *α* = 0.05 is a gain of homeostasis at target firing rate *r*_0_, and *r_i_* is a current firing rate of *i^th^* TC neuron. The current firing rate *r_i_* is computed every 120 seconds. By the end of that interval, synaptic weights are updated. The mean firing rate is estimated over 27 minutes intervals, and the simulation continues until the mean firing rate will not reach the target firing rate with 10% margins. Homeostasis requires from 11 to 22 intervals to converge to the target firing rate, ranging overall simulation time from 5 to 10 hours of model time.

The TRN inhibitory feedback is sketched as a non-specific inhibitory loop with significant delay. The GABA current is modeled as double exponential synapse with 5 ms rise and 50 ms decay time constants and −70 mV reversal potential. Note that shunting inhibition shows a much weaker effect than hyperpolarized inhibition, therefore, in some test simulations (not shown), we used up to −90 mV reversal potential to justify results reported here.

Finally, cortical excitatory feedback is modeled as a non-specific excitatory loop with significant delay. NMDA and AMPA currents have the same parameters as in rGC projections, except presynaptic single spike depression (*u*_0_), which was set to 0.7 to match much stronger paired-pulse depression in cortical axons (WG unpublished data). For the complete connectivity models, the delay for TRN connections was strictly longer than for cortical connections because there are no dLGN→TRN connections at this age, and excitation should pass the cortex before reaching TRN (Figures 3A and 4A).

### Cortical Network Model

We implement Wu et al 2020 (***Wu et al., 2020***) model as a Python-3 script for Brian2 simulator (***Stimberg et al., 2019***). Equations for excitatory neurons are the following:

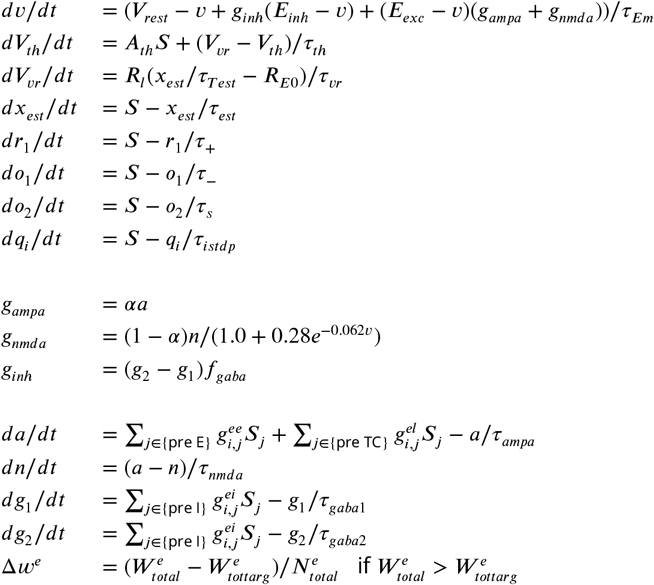

where, *v* is a membrane potential, *S* = ∑*δ*(*t*–*t*′) is a postsynaptic firing rate and *t*′ is time moment of spikes, *S_j_* are presynaptic firing rates, 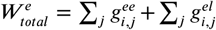 total sum of all excitatory conductance for a given neuron, 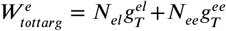 is target of total synaptic conductance for a given neuron, 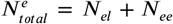 total number of excitatory synapses, *f_gaba_* is normalized coefficient scaling peak unitary inhibitory conductance to 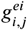.

Equations for inhibitory neurons are the following:

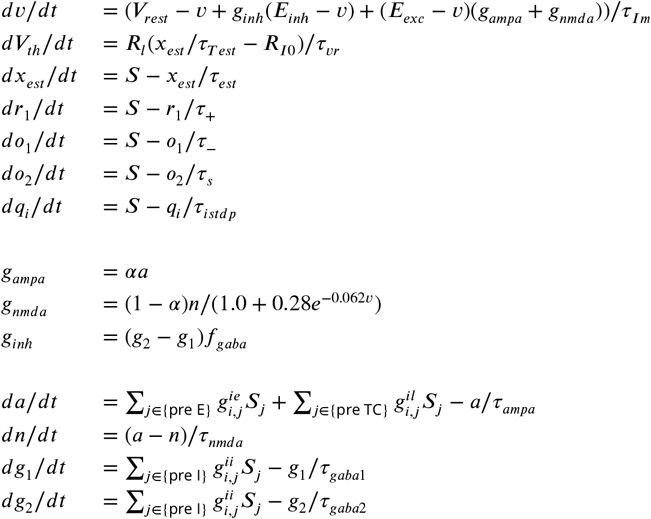

For both excitatory and inhibitory neurons, the spike is triggered when the membrane potential reaches the threshold. Subsequently, the membrane potential is reset to *V_res_*, (*v* ≥ *V_th_* : *v* → *V_rest_*) and the neuron is disabled during the refractory.

Equations for network connections and plasticity are the following:

For each E->E connection

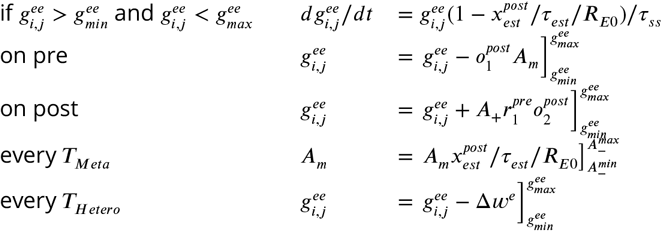
For each TC->E connection

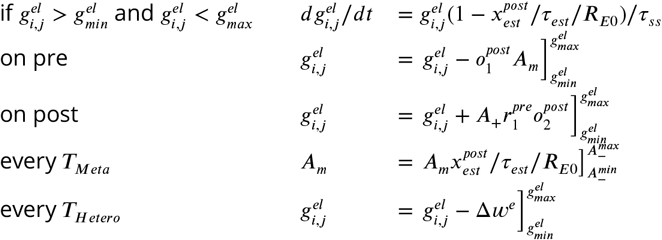
For each I->E connection

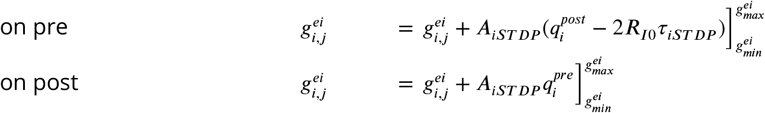

The full list of the model parameters are given in the Neuron, synaptic and plasticity parameters and Network parameters tables.

### Quantification of mutual information

To compute mutual information, we first filtered burst onset time, as in the original paper ***Butts and Rokhsar*** (***2001***), but with a shorted threshold of 0.1 second for burst onset, as TC neurons have higher firing rates than rGC neurons at P7-P10 (***Murata and Colonnese, 2018***). We then compute distributions of interburst intervals for each pear of neurons in our model, and reconstruct conditional distribution *p*(Δ*t*|*r*). Note that we will refer to both interburst intervals and interspike intervals as ISI because the results for this analysis are almost identical, but computations with interburst intervals can be performed on a regular computer with 64Gb memory, while computations with interspike intervals need more than 128Gb memory and were performed on high-memory nodes of the High-Performance Computing Cluster at the Georg Washington University (***Computing, 2020***). We then use the Shannon Mutual Information (MI), a quantitative measure of the interdependence of neuron separation (*r*) and ISI (Δ*t*), using of the conditional distributions *p*(Δ*t*|*r*) as follows:

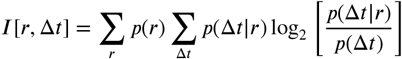

where *p*(*r*) is the prior distribution, representing the probability that two neurons chosen at random are in distance *r* apart, and 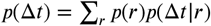 overall distribution of ISIs.

### Simulation software and Model availability

The models of individual TC neurons for EMO and the entire dLGN model were implemented as a Python-3 scripts for running the NEURON simulator (***Hines and Carnevale, 2001***). The model of the cortical network was implemented as a Python-3 script for running Bian2 simulator (***Stimberg et al., 2019***). For both network models, simulations were speeded up by employing multithreading through the ParallelContext in NEURON and through the standalone, OpenMP code generator in Brian2. Python-3 code for the optimization pipeline and both network models will be made publicly available via the ModelDB website (***McDougal et al., 2017***) after publication of this article: http://modeldb.yale.edu/267589 (code for reviewers **xGjD5lfr3ek3dhZ**).

## Acknowledgments

This work was supported by NIH grants R01EY022730 and R01NS106244 to MTC, and EY012716 to WG. This work was completed in part with resources provided by the High Performance Computing Cluster at The George Washington University, Information Technology, Research Technology Services (***Computing, 2020***) and Neuroscience Gateway Portal (***Carnevale et al., 2014***).

---

**Table 2.** Neuron, synaptic and plasticity parameters

| Parameter | CLI parameter | value | Comment |
| --- | --- | --- | --- |
| $V_{rest}$ | /aLIF/Vrest | -62 mV | Resting membrane potential [Colonesse 2016] |
| $\tau_{Em}$ | /aLIF/Tmexc | 20 ms | Membrane time constant of excitatory neuron |
| $\tau_{Im}$ | /aLIF/Tminh | 10 ms | Membrane time constant of inhibitory neuron |
|  | /aLIF/Tref | 5 ms | Duration of refractory period |
| $\tau_{th}$ | /aLIF/exc/Ta | 100 ms | fast spike-rate threshold adaptation in excitatory neuron |
| $A_{th}$ | /aLIF/exc/Aa | 5 mV | gain of fast spike-rate threshold adaptation in excitatory neuron |
| $E_{exc}$ | /syn/Eexc | 0 mV | Reversal potential for excitatory synapses |
| $E_{inh}$ | /syn/Einh | -62 mV | Reversal potential for inhibitory synapses |
| $\tau_{ampa}$ | /syn/t2_ampa | 5 ms | AMPA decay time constant |
| $\tau_{nmda}$ | /syn/t2_nmda | 100 ms | NMDA decay time constant |
| $\alpha$ | /syn/alpha | 0.5 | NMDA to AMPA receptor weighting factor |
| $\tau_{gaba1}$ | /syn/t1_gaba | 5 ms | GABA rise time constant |
| $\tau_{gaba2}$ | /syn/t2_gaba | 50 ms | GABA decay time constant |
| $R_{E0}$ | /plast/Re0 | 5 Hz | Target firing rate of excitatory neurons |
| $R_{I0}$ | /plast/Ri0 | 13 Hz | Target firing rate of inhibitory neurons |
| $\tau_+$ | /plast/T+ | 16.8 ms | Time constant of presynaptic detector |
| $\tau_-$ | /plast/T- | 33.7 ms | Time constant of faster postsynaptic detector |
| $\tau_s$ | /plast/Ts | 114 ms | Time constant of slower postsynaptic detector |
| $A_-$ | /plast/A- | 0.0071 | Amplitude of LTD |
| $A_-^{min}$ | | 0.15 $A_-$ | Miminal amplitude of LTD |
| $A_-^{max}$ | | 1.8 $A_-$ | Maximal a mplitude of LTD |
| $A_+$ | /plast/A+ | 0.0065 | Amplitude of LTP |
| $\tau_{istdp}$ | /plast/Tistdp | 0.02 s | Time constant of synaptic trace for iSTDP |
| $A_{iSTDP}$ | /plast/Aistdp | 1 | Learning rate of iSTDP |
| $\tau_{est}$ | /plast/Test | 20 s | Time constant of firing rate estimator |
| $\tau_{ss}$ | /plast/Tss | 200 s | Time constant of synaptic scaling |
| $\tau_{vr}$ | /plast/Tvr | 200 s | Time constant of intrinsic plasticity |
| $R_l$ | /plast/RI | 0.00025 mV/s | Learning rate of intrinsic plasticity |
| $g_{min}^{ee}$ | /plast/weemin | 0.0 | Minimal E-to-E connection weight |
| $g_{max}^{ee}$ | /plast/weemax | 1.2 | Maximal E-to-E connection weight |
| $g_{min}^{ei}$ | /plast/weimin | 0.0 | Minimal I-to-E connection weight |
| $g_{max}^{ei}$ | /plast/weimax | 6.0 | Maximal I-to-E connection weight |
| $g_{min}^{el}$ | /plast/welmin | 0.0 | Minimal LGN-to-E connection weight |
| $g_{max}^{el}$ | /plast/welmax | 1.2 | Maximal LGN-to-E connection weight |
| $T_{Hetero}$ | /plast/updates/Heterosynaptic | 1 s | Update time for Heterosynaptic plasticity |
| $T_{Meta}$ | /plast/updates/Metaplasticity | 60 s | Update time for Metaplasticity |
| $g_T^{ee}$ | /target/wee | 0.2 | Target for average E-to-E connection conductance |
| $g_T^{el}$ | /target/wel | 0.78 | Target for average TC-to-E connection conductance |

**Table 3.**
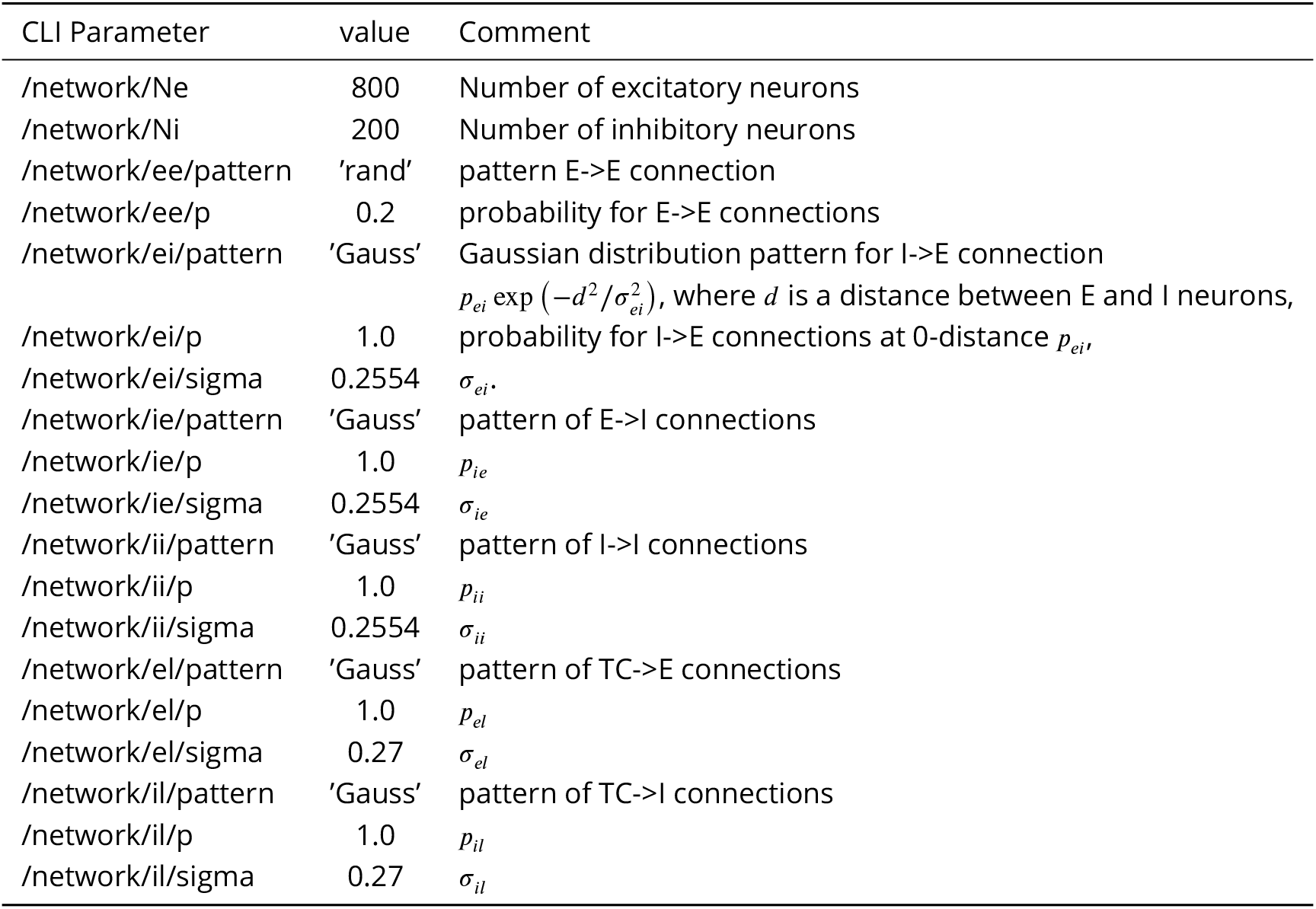
Network parameters

## Appendix 1

### Genetic algorithm with Krayzman’s adaptive weights (KAMOGA)

This EMO is built upon the algorithm for adjusting fitness (cost, objective, penalty) function weights developed by ***Eremenko et al***. (***2019***) for Reverse Monte Carlo optimization in physic. For each *i^th^* parameter set, a vector of fitness functions **F**_*i*_ = {*F_i,j_*}_*j*_ is computed, where *F_i,j_* is the *j^th^* fitness function. For each generation of the genetic algorithm (GA), **F**_*i*_ are combined into a fitness matrix **F**, and the final single-value fitness for each parameter set (*s_i_*) within the generation is obtained by multiplication of the fitness matrix by the weight vector **s** = **F** · **w**. At the begging, weights are initiated as the normalized inverse variance for each fitness function, i.e. *w_j_* = 1/*var_i_F_i,j_*, and then *w_j_* = *w_j_*/max **w**, where subscribed index (i.e., *i*) denotes the index along which variance is computed.

For each next generation, two more vectors are computed: the first is the same **s** as above, and the second is a vector of correlations between individual fitness functions within the generation and vector **s**:

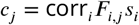

where corr_*i*_ denotes the Pearson correlation coefficient computed along *i^th^* index.

Next, iteratively, the ratio between minimal and maximal correlation is computed *r* = min **c**/ max **c**, and until this ratio is below a threshold (*θ_r_*) the algorithm performs an iteration procedure. For each iteration, the weight with minimal correlation increases, while the weight with maximal correlation decreases inversely to the number of fitness functions:

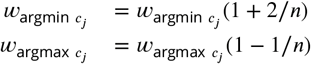

where *n* is the total number of fitness functions for MO. Then the weight vector is renormalized, and a new vector of single-value fitness for each parameter set (**s**) and a new correlation vector comprised by correlations for each fitness function (**c**) are computed as above. Iterations stop when *r* > *θ_r_*. The iteration procedure is not always converging, and *r* may systematically decrease. Therefore, there is a cap on the maximal number of iterations in which *r* decreases sequentially, and it is set to 300. If *r* decreases 300 number of iterations in a row, all weights are reset (*w_j_* = 1/*var_i_F_i,j_*) and renormalized (*w_j_* = *w_j_*/max **w**).

A critical property of KAMOGA can be seen from the point of view of the gradient descent procedure in machine learning. KAMOGA **decreases** weights for fitness functions that correlate and **increases** weights for fitness functions that anticorrelate or uncorrelate with the output. Therefore, KAMOGA is a gradient descent toward a “flatter surface” in n-dimensional space. It prevents the over- or under-representation of the individual fitness function in the weighted sum and equilibrates fitness contributions dynamically with the progression of GA. Interestingly, with *θ_r_* ≥ 0, the algorithm guarantees that all fitness functions correlate with the weighted sum.

Note that GA with elitist selection should be modified to use Krayzman’s adaptive weights. To avoid mixing fitness functions obtained with different weight vectors, the number of elites must be set to zero whenever the iteration procedure begins by condition *r* < *θ_r_*. As a result, high threshold values can force the algorithm to update weights with each generation GA, abolishing any advantages of elitism. Thus, the threshold *θ_r_* should be chosen in such a way that GA will produce at least a few generations before it loses elites. Although higher values of the threshold *θ_r_* may help to find better a balance between fitness functions, *θ_r_* ≈ 0 is a better choice for GA with elitist selection.

---

**Figure 1—figure supplement 1.**
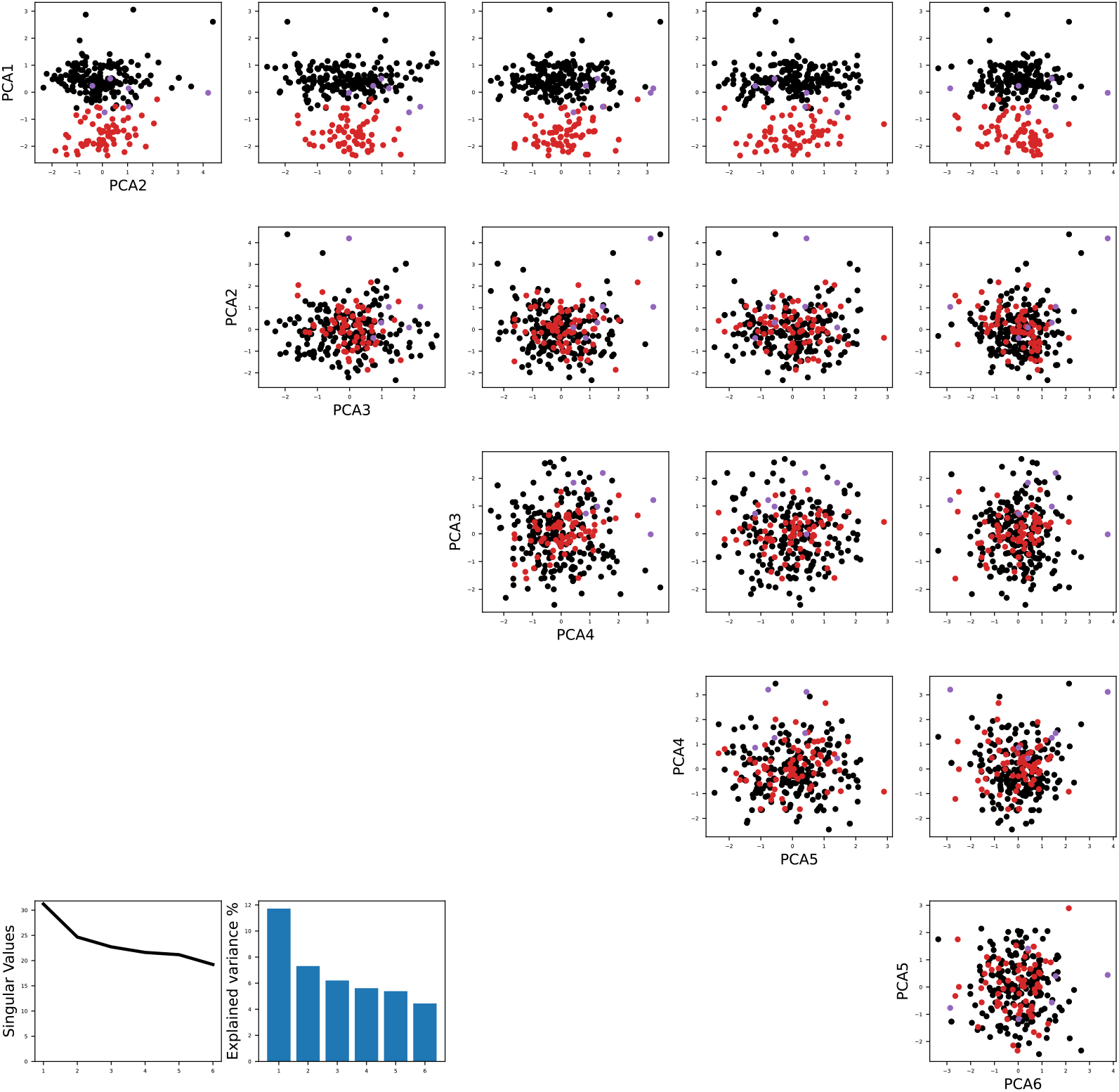
Principal component analysis shows the separation of models fitted to the same recorded neurons as in Figure 1. The color code is the same as in Figure 1B in the main text. PCA decomposition was performed using *sklearn* Python library.

**Figure 4—figure supplement 1.**
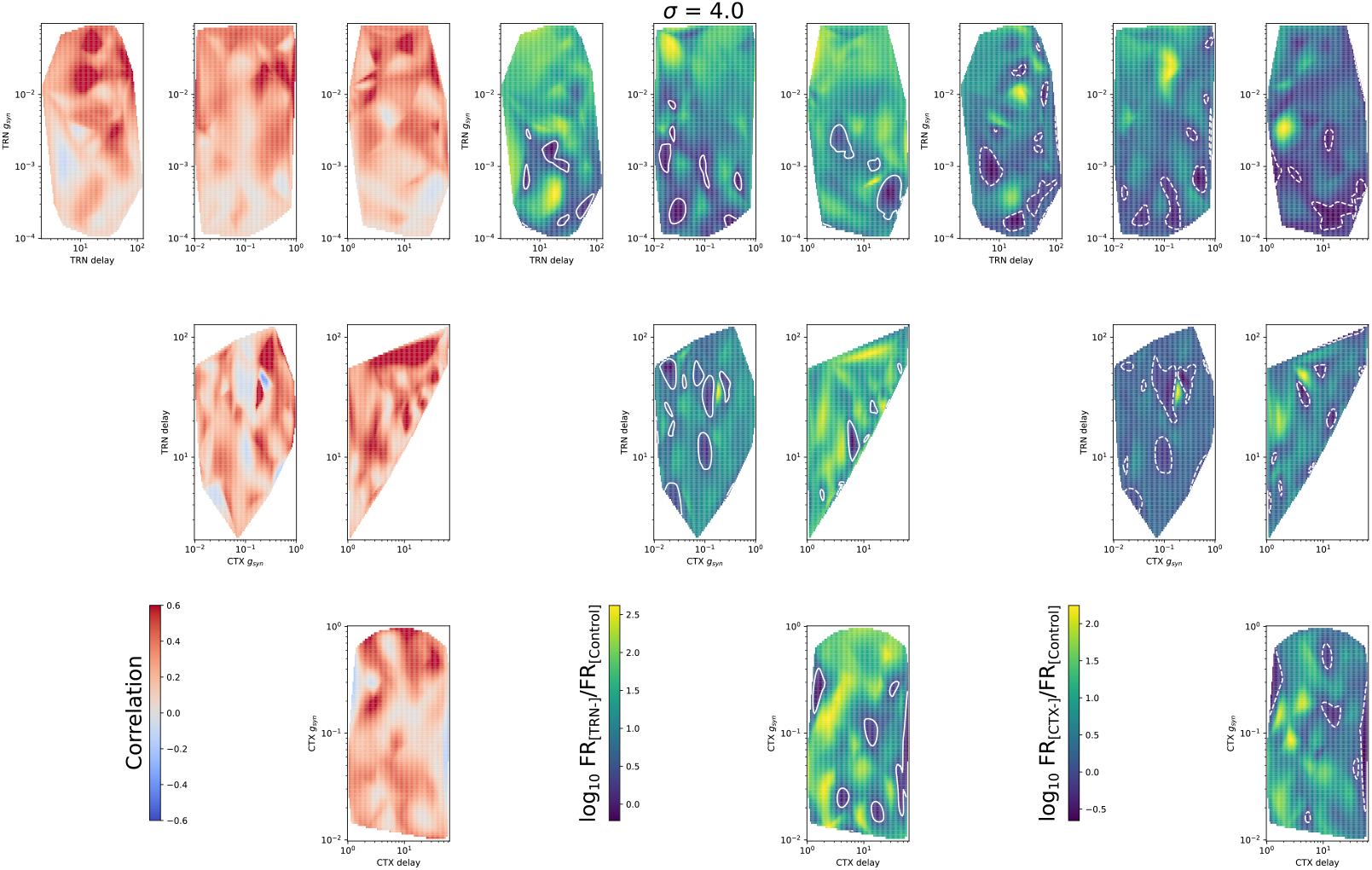
Heatmaps for mean spike correlation (left set), *F_TRN–_/F_Control_* (middle set), and *F_CTX–_ /F_Control_* (right set) for all 4 model parameters: TRN synaptic conductance *g_TAN_*, TRN delay, CTX synaptic conductance *g_CTX_*, and CTX delay. Convergence *σ* = 4.

**Figure 4—figure supplement 2.**
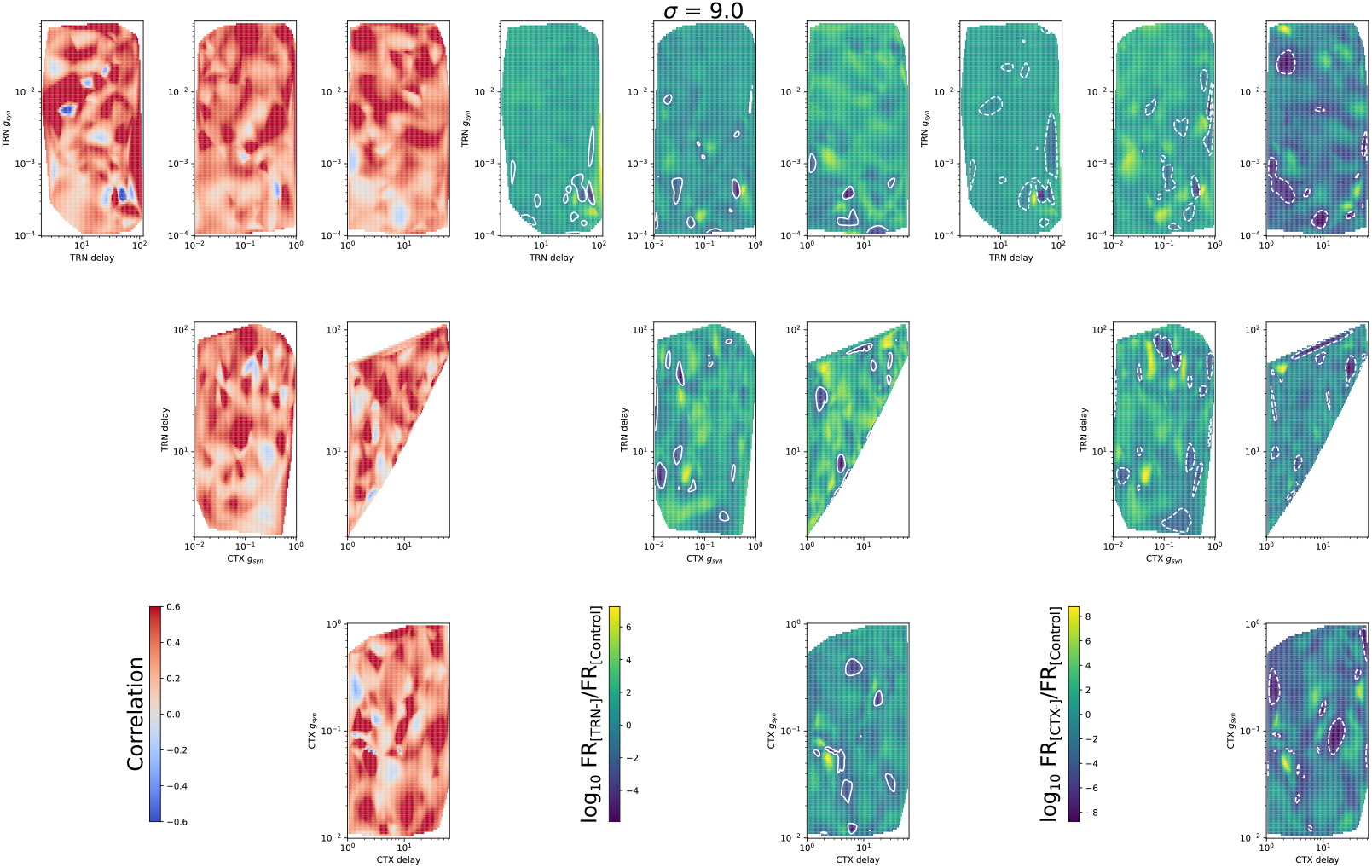
The same as in Figure 4-Figure Supplement 1, but for *σ* = 9

**Figure 6—figure supplement 1.**
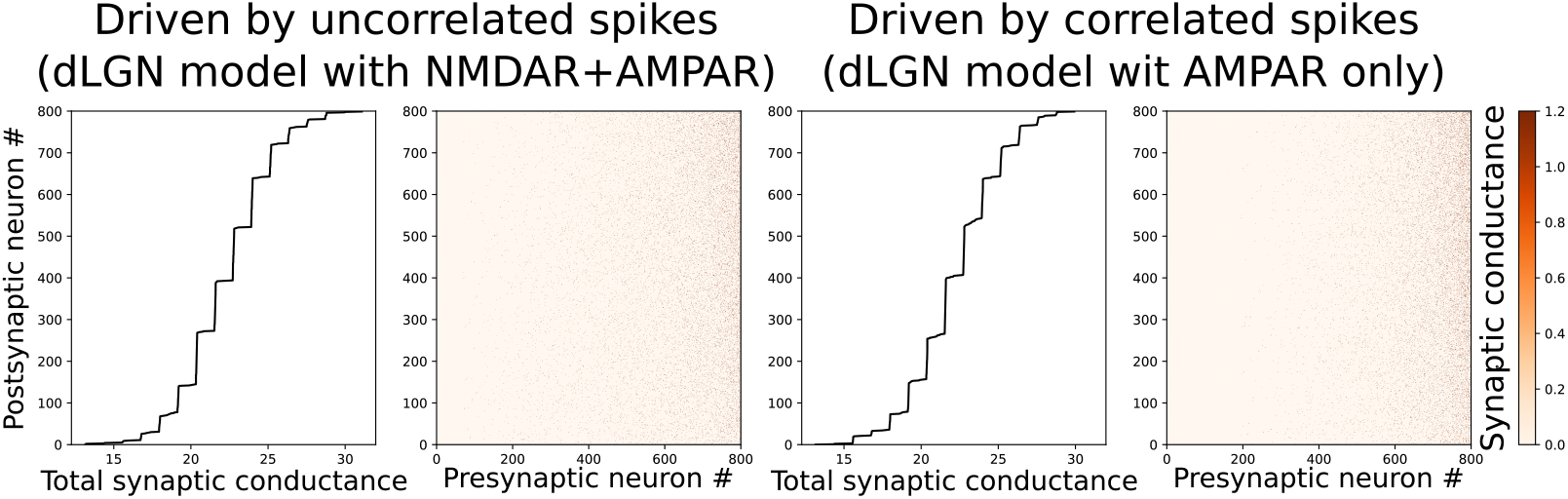
Distribution of total *input* conductance sorted from bottom up (on the left) along with matrix of excitatory-excitatory connections shown as heatmap for the same networks shown on Figure 6 A and B of the main text.

**Figure 6—figure supplement 2.**
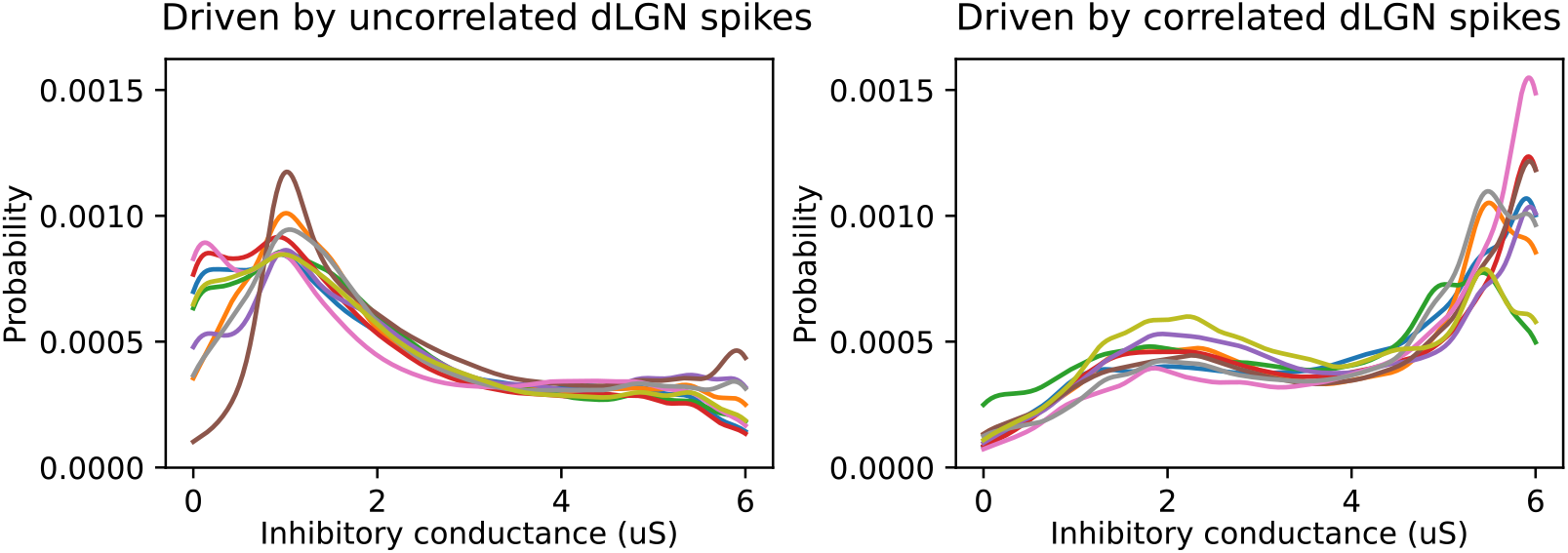
Probability I→E conducntace for 8 models driven by uncorrelated (left) and correlated (right) dLGN spikes.

## References

Almog M, Korngreen A. Is realistic neuronal modeling realistic? Journal of Neurophysiology. 2016; 116(5):2180–2209. https://doi.org/10.1152/jn.00360.2016, doi: 10.1152/jn.00360.2016, pMID: 27535372.

Arakawa H, Suzuki A, Zhao S, Tsytsarev V, Lo FS, Hayashi Y, Itohara S, Iwasato T, Erzurumlu RS. Thalamic NMDA Receptor Function Is Necessary for Patterning of the Thalamocortical Somatosensory Map and for Sensori-motor Behaviors. J Neurosci. 2014; 34(36):12001–12014. https://www.jneurosci.org/content/34/36/12001, doi: 10.1523/JNEUROSCI.1663-14.2014.

Bickford ME, Slusarczyk A, Dilger EK, Krahe TE, Kucuk C, Guido W. Synaptic development of the mouse dorsal lateral geniculate nucleus. Journal of Comparative Neurology. 2010; 518(5):622–635. https://onlinelibrary.wiley.com/doi/abs/10.1002/cne.22223, doi: 10.1002/cne.22223.

Bloomfield SA, Sherman SM. Dendritic current flow in relay cells and interneurons of the cat’s lateral geniculate nucleus. Proceedings of the National Academy of Sciences. 1989; 86(10):3911–3914. https://www.pnas.org/doi/abs/10.1073/pnas.86.10.3911, doi: 10.1073/pnas.86.10.3911.

Butts D, Kanold P. The Applicability of Spike Time Dependent Plasticity to Development. Frontiers in Synaptic Neuroscience. 2010; 2:30. https://www.frontiersin.org/article/10.3389/fnsyn.2010.00030, doi: 10.3389/fnsyn.2010.00030.

Butts DA, Feller MB, Shatz CJ, Rokhsar DS. Retinal Waves Are Governed by Collective Network Properties. The Journal of Neuroscience. 1999; 19(9):3580–3593. http://www.jneurosci.org/content/19/9/3580.abstract.

Butts DA, Kanold PO, Shatz CJ. A Burst-Based âHebbianâ Learning Rule at Retinogeniculate Synapses Links Retinal Waves to Activity-Dependent Refinement. PLoS Biol. 2007 03; 5(3):e61. http://dx.doi.org/10.1371%2Fjournal.pbio.0050061, doi: 10.1371/journal.pbio.0050061.

Butts DA, Rokhsar DS. The Information Content of Spontaneous Retinal Waves. The Journal of Neuroscience. 2001; 21 (3):961–973. http://www.jneurosci.org/content/21/3/961.abstract.

Butts DA, Weng C, Jin J, Yeh CI, Lesica NA, Alonso JM, Stanley GB. Temporal precision in the neural code and the timescales of natural vision. Nature. 2007 Sep; 449(7158):92–95. https://doi.org/10.1038/nature06105.

Carnevale T, Majumdar A, Sivagnanam S, Yoshimoto K, Astakhov V, Bandrowski A, Martone M. The neuroscience gateway portal: high performance computing made easy. BMC Neuroscience. 2014 jul; 15(S1). doi: 10.1186/1471-2202-15-s1-p101.

Charalambakis NE, Govindaiah G, Campbell PW, Guido W. Developmental Remodeling of Thalamic Interneurons Requires Retinal Signaling. Journal of Neuroscience. 2019; 39(20):3856–3866. https://www.jneurosci.org/content/39/20/3856, doi: 10.1523/JNEUROSCI.2224-18.2019.

Chen C, Blitz DM, Regehr WG. Contributions of Receptor Desensitization and Saturation to Plasticity at the Retinogeniculate Synapse. Neuron. 2002; 33(5):779 – 788. http://www.sciencedirect.com/science/article/pii/S0896627302006116, doi: https://doi.org/10.1016/S0896-6273(02)00611-6.

Chen C, Regehr WG. Developmental Remodeling of the Retinogeniculate Synapse. Neuron. 2000; 28(3):955 – 966. http://www.sciencedirect.com/science/article/pii/S0896627300001665, doi: https://doi.org/10.1016/S0896-6273(00)00166-5.

Colonnese MT. Rapid Developmental Emergence of Stable Depolarization during Wakefulness by Inhibitory Balancing of Cortical Network Excitability. Journal of Neuroscience. 2014; 34(16):5477–5485. http://www.jneurosci.org/content/34/16/5477, doi: 10.1523/JNEUROSCI.3659-13.2014.

Colonnese MT, Constantine-Paton M. Chronic NMDA Receptor Blockade from Birth Increases the Sprouting Capacity of Ipsilateral Retinocollicular Axons without Disrupting Their Early Segregation. Journal of Neuroscience. 2001; 21(5):1557–1568. https://www.jneurosci.org/content/21/5/1557, doi: 10.1523/JNEUROSCI.21-05-01557.2001.

Colonnese MT, Constantine-Paton M. Developmental period for N-methyl-D-aspartate (NMDA) receptor-dependent synapse elimination correlated with visuotopic map refinement. Journal of Comparative Neurology. 2006; 494(5):738–751. https://onlinelibrary.wiley.com/doi/abs/10.1002/cne.20841, doi: https://doi.org/10.1002/cne.20841.

Colonnese MT, Phillips MA. Thalamocortical function in developing sensory circuits. Current Opinion in Neurobiology. 2018; 52:72 – 79. http://www.sciencedirect.com/science/article/pii/S0959438817302970, doi: https://doi.org/10.1016/j.conb.2018.04.019, systems Neuroscience.

Colonnese MT, Shen J, Murata Y. Uncorrelated Neural Firing in Mouse Visual Cortex during Spontaneous Retinal Waves. Frontiers in Cellular Neuroscience. 2017; 11:289. https://www.frontiersin.org/article/10.3389/fncel.2017.00289, doi: 10.3389/fncel.2017.00289.

Colonnese MT, Shi J, Constantine-Paton M. Chronic NMDA Receptor Blockade From Birth Delays the Maturation of NMDA Currents, but Does Not Affect AMPA/Kainate Currents. Journal of Neurophysiology. 2003; 89(1):57–68. https://doi.org/10.1152/jn.00049.2002, doi: 10.1152/jn.00049.2002, pMID: 12522159.

Computing TGWUHP; 2020. https://arxiv.org/abs/2003.13629/.

Connelly WM, Crunelli V, Errington AC. Passive Synaptic Normalization and Input Synchrony-Dependent Amplification of Cortical Feedback in Thalamocortical Neuron Dendrites. Journal of Neuroscience. 2016; 36(13):3735–3754. https://www.jneurosci.org/content/36/13/3735, doi: 10.1523/JNEUROSCI.3836-15.2016.

Deb K, Pratap A, Agarwal S, Meyarivan T. A fast and elitist multiobjective genetic algorithm: NSGA-II. IEEE Transactions on Evolutionary Computation. 2002 apr; 6(2):182–197. doi: 10.1109/4235.996017.

Deb K. Multi-Objective Optimization Using Evolutionary Algorithms. Wiley; 2001.

Dilger EK, Krahe TE, Morhardt DR, Seabrook TA, Shin HS, Guido W. Absence of Plateau Potentials in dLGN Cells Leads to a Breakdown in Retinogeniculate Refinement. Journal of Neuroscience. 2015; 35(8):3652–3662. https://www.jneurosci.org/content/35/8/3652, doi: 10.1523/JNEUROSCI.2343-14.2015.

Dilger EK, Shin HS, Guido W. Requirements for synaptically evoked plateau potentials in relay cells of the dorsal lateral geniculate nucleus of the mouse. The Journal of Physiology. 2011; 589(4):919–937. https://physoc.onlinelibrary.wiley.com/doi/abs/10.1113/jphysiol.2010.202499, doi: 10.1113/jphysiol.2010.202499.

Drew PJ, Abbott LF. Extending the effects of spike-timing-dependent plasticity to behavioral timescales. Proceedings of the National Academy of Sciences. 2006 jun; 103(23):8876–8881. doi: 10.1073/pnas.0600676103.

Dura-Bernal S, Neymotin SA, Kerr CC, Sivagnanam S, Majumdar A, Francis JT, Lytton WW. Evolutionary algorithm optimization of biological learning parameters in a biomimetic neuroprosthesis. IBM Journal of Research and Development. 2017 mar;61(2/3):6:1–6:14. doi: 10.1147/jrd.2017.2656758.

Eglen SJ, Weeks M, Jessop M, Simonotto J, Jackson T, Sernagor E. A data repository and analysis framework for spontaneous neural activity recordings in developing retina. GigaScience. 2014 03; 3(1). https://doi.org/10.1186/2047-217X-3-3, doi: 10.1186/2047-217X-3-3, 2047-217X-3-3.

El-Danaf RN, Krahe TE, Dilger EK, Bickford ME, Fox MA, Guido W. Developmental remodeling of relay cells in the dorsal lateral geniculate nucleus in the absence of retinal input. Neural Development. 2015 Jul; 10(1):19. https://doi.org/10.1186/s13064-015-0046-6.

Elstrott J, Feller MB. Direction-Selective Ganglion Cells Show Symmetric Participation in Retinal Waves During Development. The Journal of Neuroscience. 2010; 30(33):11197–11201. http://www.jneurosci.org/content/30/33/11197.abstract, doi: 10.1523/JNEUROSCI.2302-10.2010.

Eremenko M, Krayzman V, Bosak A, Playford HY, Chapman KW, Woicik JC, Ravel B, Levin I. Local atomic order and hierarchical polar nanoregions in a classical relaxor ferroelectric. Nature Communications. 2019 jun; 10(1). doi: 10.1038/s41467-019-10665-4.

Evrard A, Ropert N. Early Development of the Thalamic Inhibitory Feedback Loop in the Primary Somatosensory System of the Newborn Mice. Journal of Neuroscience. 2009; 29(31):9930–9940. https://www.jneurosci.org/content/29/31/9930, doi: 10.1523/JNEUROSCI.1671-09.2009.

Ewald RC, Cline HT. In: Van Dongen AM, editor. NMDA Receptors and Brain Development. Boca Raton (FL); 2009..

Feller MB. Retinal waves are likely to instruct the formation of eye-specific retinogeniculate projections. Neural Development. 2009 Jul; 4(1):24. https://doi.org/10.1186/1749-8104-4-24.

Fiete IR, Senn W, Wang CZH, Hahnloser RHR. Spike-Time-Dependent Plasticity and Heterosynaptic Competition Organize Networks to Produce Long Scale-Free Sequences of Neural Activity. Neuron. 2010; 65(4):563–576. https://www.sciencedirect.com/science/article/pii/S0896627310000917, doi: https://doi.org/10.1016/j.neuron.2010.02.003.

Gjorgjieva J, Clopath C, Audet J, Pfister JP. A triplet spike-timing-dependent plasticity model generalizes the Bienenstock-Cooper-Munro rule to higher-order spatiotemporal correlations. Proceedings of the National Academy of Sciences. 2011; 108(48):19383–19388. https://www.pnas.org/doi/abs/10.1073/pnas.1105933108, doi: 10.1073/pnas.1105933108.

Guido W. Development, form, and function of the mouse visual thalamus. Journal of Neurophysiology. 2018; 120(1):211–225. https://doi.org/10.1152/jn.00651.2017, doi: 10.1152/jn.00651.2017, pMID: 29641300.

Hahm JO, Langdon RB, Sur M. Disruption of retinogeniculate afferent segregation by antagonists to NMDA receptors. Nature. 1991 jun; 351(6327):568–570. doi: 10.1038/351568a0.

Hauser JL, Liu X, Litvina EY, Chen C. Prolonged synaptic currents increase relay neuron firing at the developing retinogeniculate synapse. Journal of Neurophysiology. 2014; 112(7):1714–1728. https://doi.org/10.1152/jn.00451.2014, doi: 10.1152/jn.00451.2014, pMID: 24966302.

Hines ML, Carnevale NT. Neuron: A Tool for Neuroscientists. The Neuroscientist. 2001; 7(2):123–135. https://doi.org/10.1177/107385840100700207, doi: 10.1177/107385840100700207, pMID: 11496923.

Hooks BM, Chen C. Circuitry Underlying Experience-Dependent Plasticity in the Mouse Visual System. Neuron. 2020; 106(1):21–36. https://www.sciencedirect.com/science/article/pii/S089662732030057X, doi: https://doi.org/10.1016/j.neuron.2020.01.031.

Huang L, Pallas SL. NMDA Antagonists in the Superior Colliculus Prevent Developmental Plasticity But Not Visual Transmission or Map Compression. Journal of Neurophysiology. 2001; 86(3):1179–1194. https://doi.org/10.1152/jn.2001.86.3.1179, doi: 10.1152/jn.2001.86.3.1179, pMID: 11535668.

Huberman AD, Feller MB, Chapman B. Mechanisms Underlying Development of Visual Maps and Receptive Fields. Annual Review of Neuroscience. 2008; 31(1):479–509. https://doi.org/10.1146/annurev.neuro.31.060407.125533, doi: 10.1146/annurev.neuro.31.060407.125533, pMID: 18558864.

Iavarone E, Yi J, Shi Y, Zandt BJ, O’Reilly C, Van Geit W, Rössert C, Markram H, Hill SL. Experimentally-constrained biophysical models of tonic and burst firing modes in thalamocortical neurons. PLOS Computational Biology. 2019 05; 15(5):1–23. https://doi.org/10.1371/journal.pcbi.1006753, doi: 10.1371/journal.pcbi.1006753.

Iwasato T, Datwani A, Wolf AM, Nishiyama H, Taguchi Y, Tonegawa S, Knöpfel T, Erzurumlu RS, Itohara S. Cortex-restricted disruption of NMDAR1 impairs neuronal patterns in the barrel cortex. Nature. 2000 aug; 406(6797):726–731. doi: 10.1038/35021059.

Jacobsen RB, Ulrich D, Huguenard JR. GABAB and NMDA Receptors Contribute to Spindle-Like Oscillations in Rat Thalamus In Vitro. Journal of Neurophysiology. 2001; 86(3):1365–1375. https://doi.org/10.1152/jn.2001.86.3.1365, doi: 10.1152/jn.2001.86.3.1365, pMID: 11535683.

Jaubert-Miazza L, Green E, Lo FS, Bui K, Mills J, Guido W. Structural and functional composition of the developing retinogeniculate pathway in the mouse. Visual Neuroscience. 2005; 22(5):661–676. doi: 10.1017/S0952523805225154.

Kano M, Hashimoto K. Synapse elimination in the central nervous system. Current Opinion in Neurobiology. 2009 apr; 19(2):154–161. doi: 10.1016/j.conb.2009.05.002.

Katz LC, Shatz CJ. Synaptic Activity and the Construction of Cortical Circuits. Science. 1996 nov; 274(5290):1133–1138. doi: 10.1126/science.274.5290.1133.

Kesner P, Schohl A, Warren EC, Ma F, Ruthazer ES. Postsynaptic and Presynaptic NMDARs Have Distinct Roles in Visual Circuit Development. Cell Reports. 2020; 32(4):107955. https://www.sciencedirect.com/science/article/pii/S2211124720309360, doi: https://doi.org/10.1016/j.celrep.2020.107955.

Kirkby L, Sack G, Firl A, Feller M. A Role for Correlated Spontaneous Activity in the Assembly of Neural Circuits. Neuron. 2013; 80(5):1129 – 1144. http://www.sciencedirect.com/science/article/pii/S0896627313009343, doi: https://doi.org/10.1016/j.neuron.2013.10.030.

Kirmse K, Zhang C. Principles of GABAergic signaling in developing cortical network dynamics. Cell Reports. 2022; 38(13):110568. https://www.sciencedirect.com/science/article/pii/S2211124722003126, doi: https://doi.org/10.1016/j.celrep.2022.110568.

Kuo MC, Dringenberg HC. Comparison of long-term potentiation (LTP) in the medial (monocular) and lateral (binocular) rat primary visual cortex. Brain research. 2012 Dec; 1488:51–9.

Lee H, Brott BK, Kirkby LA, Adelson JD, Cheng S, Feller MB, Datwani A, Shatz CJ. Synapse elimination and learning rules co-regulated by MHC class I H2-Db. Nature. 2014 mar; 509(7499):195–200. doi: 10.1038/nature13154.

Liang L, Chen C. Organization, Function, and Development of the Mouse Retinogeniculate Synapse. Annual Review of Vision Science. 2020 sep; 6(1):261–285. doi: 10.1146/annurev-vision-121219-081753.

Litwin-Kumar A, Doiron B. Slow dynamics and high variability in balanced cortical networks with clustered connections. Nature neuroscience. 2012; 15(11):1498–1505.

Litwin-Kumar A, Doiron B. Formation and maintenance of neuronal assemblies through synaptic plasticity. Nature communications. 2014; 5.

Liu X, Chen C. Different Roles for AMPA and NMDA Receptors in Transmission at the Immature Retinogeniculate Synapse. Journal of Neurophysiology. 2008; 99(2):629–643. https://doi.org/10.1152/jn.01171.2007, doi: 10.1152/jn.01171.2007, pMID: 18032559.

Lo FS, Ziburkus J, Guido W. Synaptic Mechanisms Regulating the Activation of a Ca2+-Mediated Plateau Potential in Developing Relay Cells of the LGN. Journal of Neurophysiology. 2002; 87(3):1175–1185. https://doi.org/10.1152/jn.00715.1999, doi: 10.1152/jn.00715.1999, pMID: 11877491.

Maccione A, Hennig MH, Gandolfo M, Muthmann O, Coppenhagen J, Eglen SJ, Berdondini L, Sernagor E. Following the ontogeny of retinal waves: pan-retinal recordings of population dynamics in the neonatal mouse. The Journal of physiology. 2014; 592(7):1545–1563.

Marder E. Variability, compensation, and modulation in neurons and circuits. Proceedings of the National Academy of Sciences. 2011; 108(Supplement 3):15542–15548. https://www.pnas.org/content/108/Supplement_3/15542, doi: 10.1073/pnas.1010674108.

McDougal RA, Morse TM, Carnevale T, Marenco L, Wang R, Migliore M, Miller PL, Shepherd GM, Hines ML. Twenty years of ModelDB and beyond: building essential modeling tools for the future of neuroscience. Journal of Computational Neuroscience. 2017 Feb; 42(1):1–10. https://doi.org/10.1007/s10827-016-0623-7.

Minlebaev M, Colonnese M, Tsintsadze T, Sirota A, Khazipov R. Early Gamma Oscillations Synchronize Developing Thalamus and Cortex. Science. 2011; 334(6053):226–229. http://science.sciencemag.org/content/334/6053/226, doi: 10.1126/science.1210574.

Mizuno H, Rao MS, Mizuno H, Sato T, Nakazawa S, Iwasato T. NMDA Receptor Enhances Correlation of Spontaneous Activity in Neonatal Barrel Cortex. Journal of Neuroscience. 2021; 41(6):1207–1217. https://www.jneurosci.org/content/41/6/1207, doi: 10.1523/JNEUROSCI.0527-20.2020.

Murata Y, Colonnese MT. An excitatory cortical feedback loop gates retinal wave transmission in rodent thalamus. eLife. 2016 oct; 5:e18816. https://dx.doi.org/10.7554/eLife.18816, doi: 10.7554/eLife.18816.

Murata Y, Colonnese MT. Thalamus Controls Development and Expression of Arousal States in Visual Cortex. Journal of Neuroscience. 2018; 38(41):8772–8786. https://www.jneurosci.org/content/38/41/8772, doi: 10.1523/JNEUROSCI.1519-18.2018.

Murata Y, Colonnese MT. Thalamic inhibitory circuits and network activity development. Brain Research. 2019; 1706:13 – 23. http://www.sciencedirect.com/science/article/pii/S0006899318305365, doi: https://doi.org/10.1016/j.brainres.2018.10.024.

Neymotin SA, Suter BA, Dura-Bernal S, Shepherd GMG, Migliore M, Lytton WW. Optimizing computer models of corticospinal neurons to replicate in vitro dynamics. Journal of Neurophysiology. 2017; 117(1):148–162. https://doi.org/10.1152/jn.00570.2016, doi: 10.1152/jn.00570.2016, pMID: 27760819.

Niell CM, Scanziani M. How Cortical Circuits Implement Cortical Computations: Mouse Visual Cortex as a Model. Annual Review of Neuroscience. 2021; 44(1):517–546. https://doi.org/10.1146/annurev-neuro-102320-085825, doi: 10.1146/annurev-neuro-102320-085825, pMID: 33914591.

Pinault D. The thalamic reticular nucleus: structure, function and concept. Brain Research Reviews. 2004 aug; 46(1):1–31. doi: 10.1016/j.brainresrev.2004.04.008.

Pinsky PF, Rinzel J. Intrinsic and network rhythmogenesis in a reduced traub model for CA3 neurons. Journal of Computational Neuroscience. 1994; 1(1):39–60. http://dx.doi.org/10.1007/BF00962717, doi: 10.1007/BF00962717.

Prinz AA, Billimoria CP, Marder E. Alternative to Hand-Tuning Conductance-Based Models: Construction and Analysis of Databases of Model Neurons. Journal of Neurophysiology. 2003; 90(6):3998–4015. https://doi.org/10.1152/jn.00641.2003, doi: 10.1152/jn.00641.2003, pMID: 12944532.

Prinz AA, Bucher D, Marder E. Similar network activity from disparate circuit parameters. Nature Neuroscience. 2004 Dec; 7(12):1345–1352. https://doi.org/10.1038/nn1352.

Riyahi P, Phillips MA, Colonnese MT. Input-Independent Homeostasis of Developing Thalamocortical Activity. eneuro. 2021 may; 8(3):ENEURO.0184-21.2021. doi: 10.1523/eneuro.0184-21.2021.

Rocha M, Sur M. Rapid acquisition of dendritic spines by visual thalamic neurons after blockade of N-methyl-D-aspartate receptors. Proceedings of the National Academy of Sciences. 1995; 92(17):8026–8030. https://www.pnas.org/doi/abs/10.1073/pnas.92.17.8026, doi: 10.1073/pnas.92.17.8026.

Rochefort NL, Garaschuk O, Milos RI, Narushima M, Marandi N, Pichler B, Kovalchuk Y, Konnerth A. Sparsification of neuronal activity in the visual cortex at eye-opening. Proceedings of the National Academy of Sciences. 2009; 106(35):15049–15054. https://www.pnas.org/doi/abs/10.1073/pnas.0907660106, doi: 10.1073/pnas.0907660106.

Rumpel S, Hatt H, Gottmann K. Silent Synapses in the Developing Rat Visual Cortex: Evidence for Postsynaptic Expression of Synaptic Plasticity. Journal of Neuroscience. 1998; 18(21):8863–8874. https://www.jneurosci.org/content/18/21/8863, doi: 10.1523/JNEUROSCI.18-21-08863.1998.

Sailamul P, Jang J, Paik SB. Synaptic convergence regulates synchronization-dependent spike transfer in feedforward neural networks. Journal of Computational Neuroscience. 2017; 43(3):189–202. https://doi.org/10.1007/s10827-017-0657-5, doi: 10.1007/s10827-017-0657-5.

Sanes JR, Lichtman JW. DEVELOPMENT OF THE VERTEBRATE NEUROMUSCULAR JUNCTION. Annual Review of Neuroscience. 1999 mar; 22(1):389–442. doi: 10.1146/annurev.neuro.22.1.389.

Seabrook TA, Burbridge TJ, Crair MC, Huberman AD. Architecture, Function, and Assembly of the Mouse Visual System. Annual Review of Neuroscience. 2017; 40(1):499–538. https://doi.org/10.1146/annurev-neuro-071714-033842, doi: 10.1146/annurev-neuro-071714-033842, pMID: 28772103.

Shah RD, Crair MC. Retinocollicular Synapse Maturation and Plasticity Are Regulated by Correlated Retinal Waves. The Journal of Neuroscience. 2008; 28(1):292–303. http://www.jneurosci.org/content/28/1/292.abstract, doi: 10.1523/JNEUROSCI.4276-07.2008.

Sherman SM, Guillery RW. 311Thalamus. In: The Synaptic Organization of the Brain Oxford University Press; 2004.https://doi.org/10.1093/acprof:oso/9780195159561.003.0008, doi: 10.1093/acprof:oso/9780195159561.003.0008.

Siegel F, Heimel J Peters J, Lohmann C. Peripheral and Central Inputs Shape Network Dynamics in the Developing Visual Cortex In Vivo. Current Biology. 2012; 22(3):253 – 258. http://www.sciencedirect.com/science/article/pii/S0960982211014035, doi: https://doi.org/10.1016/j.cub.2011.12.026.

Stafford BK, Sher A, Litke AM, Feldheim DA. Spatial-temporal patterns of retinal waves underlying activity-dependent refinement of retinofugal projections. Neuron. 2009; 64(2):200–212.

Stent GS. A Physiological Mechanism for Hebb’s Postulate of Learning. Proceedings of the National Academy of Sciences. 1973; 70(4):997–1001. https://www.pnas.org/doi/abs/10.1073/pnas.70.4.997, doi: 10.1073/pnas.70.4.997.

Stimberg M, Brette R, Goodman DF. Brian 2, an intuitive and efficient neural simulator. eLife. 2019 aug; 8:e47314. https://doi.org/10.7554/eLife.47314, doi: 10.7554/eLife.47314.

Thompson A, Gribizis A, Chen C, Crair MC. Activity-dependent development of visual receptive fields. Current Opinion in Neurobiology. 2017; 42:136–143. https://www.sciencedirect.com/science/article/pii/S0959438816302604, doi: https://doi.org/10.1016/j.conb.2016.12.007, developmental neuroscience.

Tonda A. Inspyred: Bio-inspired algorithms in Python. Genetic Programming and Evolvable Machines. 2020 Jun; 21(1):269–272. https://doi.org/10.1007/s10710-019-09367-z.

Tsodyks t, Uziel A, Markram H. Synchrony Generation in Recurrent Networks with Frequency-Dependent Synapses. Journal of Neuroscience. 2000; 20(1):RC50–RC50. https://www.jneurosci.org/content/20/1/RC50, doi: 10.1523/JNEUROSCI.20-01-j0003.2000.

Usrey WM, Reid RC. Synchronous Activity In The Visual System. Annual Review of Physiology. 1999; 61(1):435–456. https://doi.org/10.1146/annurev.physiol.61.1.435, doi: 10.1146/annurev.physiol.61.1.435, pMID: 10099696.

Vogels TP, Sprekeler H, Zenke F, Clopath C, Gerstner W. Inhibitory Plasticity Balances Excitation and Inhibition in Sensory Pathways and Memory Networks. Science. 2011; 334(6062):1569–1573. https://science.sciencemag.org/content/334/6062/1569, doi: 10.1126/science.1211095.

Vonhoff F, Keshishian H. Activity-Dependent Synaptic Refinement: New Insights from Drosophila. Frontiers in Systems Neuroscience. 2017 apr; 11. doi: 10.3389/fnsys.2017.00023.

Wong ROL, Meister M, Shatz CJ. Transient period of correlated bursting activity during development of the mammalian retina. Neuron. 1993 Nov; 11(5):923–938. https://doi.org/10.1016/0896-6273(93)90122-8, doi: 10.1016/0896-6273(93)90122-8.

Wong ROL. RETINAL WAVES AND VISUAL SYSTEM DEVELOPMENT. Annual Review of Neuroscience. 1999 mar; 22(1):29–47. doi: 10.1146/annurev.neuro.22.1.29.

Wu YK, Hengen KB, Turrigiano GG, Gjorgjieva J. Homeostatic mechanisms regulate distinct aspects of cortical circuit dynamics. Proceedings of the National Academy of Sciences. 2020; 117(39):24514–24525. https://www.pnas.org/content/117/39/24514, doi: 10.1073/pnas.1918368117.

Zenke F, Gerstner W. Hebbian plasticity requires compensatory processes on multiple timescales. Philosophical Transactions of the Royal Society B: Biological Sciences. 2017 mar; 372(1715):20160259. doi: 10.1098/rstb.2016.0259.

Zhu JJ, Malinow R. Acute versus chronic NMDA receptor blockade and synaptic AMPA receptor delivery. Nature Neuroscience. 2002 apr; 5(6):513–514. doi: 10.1038/nn0602-850.

